# dynamAedes: a unified modelling framework for invasive *Aedes* mosquitoes

**DOI:** 10.1101/2021.12.21.473628

**Authors:** Daniele Da Re, Wim Van Bortel, Friederike Reuss, Ruth Müller, Sebastien Boyer, Fabrizio Montarsi, Silvia Ciocchetta, Daniele Arnoldi, Giovanni Marini, Annapaola Rizzoli, Gregory L’Ambert, Guillaume Lacour, Constantianus J.M. Koenraadt, Sophie O. Vanwambeke, Matteo Marcantonio

## Abstract

1. Mosquito species belonging to the genus *Aedes* have attracted the interest of scientists and public health officers for their invasive species traits and efficient capacity of transmitting viruses affecting humans. Some of these species were brought outside their native range by human activities such as trade and tourism, and colonised new regions thanks to a unique combination of eco-physiological traits.
2. Considering mosquito physiological and behavioural traits to understand and predict the spatial and temporal population dynamics is thus a crucial step to develop strategies to mitigate the local densities of invasive *Aedes* populations.
3. Here, we synthesised the life cycle of four invasive *Aedes* species (*Ae. aegypti*, *Ae. albopictus*, *Ae. japonicus* and *Ae. koreicus*) in a single multi-scale stochastic modelling framework which we coded in the R package `dynamAedes`. We designed a stage-based and time-discrete stochastic model driven by temperature, photo-period and inter-specific larval competition that can be applied to three different spatial scales: punctual, local and regional. These spatial scales consider different degrees of spatial complexity and data availability, by accounting for both active and passive dispersal of mosquito species as well as for the heterogeneity of the input temperature data.
4. Our overarching aim was to provide a flexible, open-source and user-friendly tool rooted in the most updated knowledge on species biology which could be applied to the management of invasive *Aedes* populations as well as for more theoretical ecological inquiries.

## 1 Introduction

Some mosquito species within the *Aedes* taxon have a unique combination of biological traits such as: 1) efficient transmission of viruses debilitating for humans and animals (Gratz, 2004; Hurk et al., 2011; Souza-Neto et al., 2019), 2) eco-physiological plasticity that allows for rapid adaptation (Kramer et al., 2021) and exploitation of novel environments created by humans (McBride et al., 2014a), 3) egg stage with high resistance to dry and cold conditions which facilitate displacements over long ecological and geographical distances (Thomas et al., 2012; Versteirt et al., 2012; Kaufman and Fonseca, 2014). Some of these species were accidentally brought outside their native areas by human activities and colonised new regions thanks to a unique combination of eco-physiological traits. These mosquitoes, often referred as “*Aedes* invasive mosquitoes” (AIM), have attracted the interest of scientists and public health officers and much effort has been done to unravel their physiological and behavioural traits. Among these species, *Ae. aegypti*, *Ae. albopictus*, *Ae. japonicus* and *Ae. koreicus* showed a rapid expansion of their geographical range, with the first two species often causing an important burden on public health. As a consequence, large experimental and observational datasets on the relationship between water or air temperature and physiological parameters have been collected and used to develop mechanistic models that reproduce the basic life cycle of these four species (e.g., for *Ae. aegypti* Focks et al., 1993a,b; Otero et al., 2006; Da Re et al., 2021; Caldwell et al., 2021; for *Ae. albopictus* Tran et al., 2013; Erguler et al., 2016; Metelmann et al., 2019; Pasquali et al., 2020; Tran et al., 2020; for *Ae. japonicus* Wieser et al., 2019; for *Ae. koreicus* Marini et al., 2019b). The inclusion of such functions, which describe physiological and developmental rates, into modelling frameworks allow for more reliable model extrapolations, as the chances of biological unrealistic outcome may be lower compared to pure correlative model approaches (Kearney, 2006; Kearney and Porter, 2009). Comparisons between modelled and observed population trends showed that such mechanistic models can be used, for example, to understand population dynamics in space and time, and thus can enhance the pest control strategies against AIM (Baldacchino et al., 2015).

Models targeting AIM developed so far aimed to simulate the population dynamics of only one species at the time and for a single qualitative (i.e. “individual”, “container” or “household”) or quantitative (cell in a lattice grid) spatial scale. Moreover, only a few of these models have been made readily operational, for example by organising them in open-access, user-friendly software or libraries with sufficient documentation for practical applications (Stallman, 1985). SkeeterBuster, a container-level population dynamical model for *Ae. aegypti*, has been the first agent-based model for mosquitoes made available as a free (but not open-source) software (Magori et al., 2009). Concerning *Ae. albopictus*, Erguler et al. (2016) made available a model developed as a Python library that was after-wards wrapped into the R package `albopictus`. More recently, the European Centre for Disease Control (ECDC) has provided a free and open-source adaptation of a model initially developed by Tran et al. (2013), making it accessible via the R Shiny application `AedesRisk`^1^. Similarly, two generic age and stage-structured discrete-time population dynamics models, also applicable to mosquitoes, were proposed in the last few years: `stagePop` (Kettle and Nutter 2015; applied to *Ae. japonicus* in Wieser et al. 2019 ) and `sPop` (Erguler, 2018). Despite the current availability of models applicable to invasive *Aedes*, none of them can be directly generalised while retaining biological credibility, e.g. species-specific models work only for a single species whereas generic models may over-simplify the life cycle structure or are not equipped with species-specific physiological parameters. Hence, if users decide to use generic models, they need to screen the scientific literature, filter and manipulate experimental data (often scarce and non-standardised) to inform models on the species of interest. Moreover, often models do not consider mosquito dispersal or even completely lack spatial structure.

Here, we synthesised the life cycle of four AIM species: *Ae. aegypti*, *Ae. albopictus*, *Ae. japonicus* and *Ae. koreicus* in a single modelling framework, which we coded in the R package `dynamAedes`. We designed a stage-based and time discrete stochastic model informed by temperature and photoperiod that can be applied to three different spatial scales: punctual, local and regional. These spatial scales were thought to meet different degrees of spatial complexity and data availability, by accounting for both active and human-mediated passive dispersal of the modelled mosquito species as well as for the heterogeneity of temperature data. Our overarching aim was to provide a flexible and open-source tool which could be used for applications related to the management of AIM populations but also for more theoretical ecological inquiries. We described and assessed the model using observational mosquito data and then showed how to use the R package with coding examples and relevant case studies.

## 2 Materials and methods

### 2.1 A summary of invasive *Aedes* species ecology

#### 2.1.1 Aedes aegypti

*Aedes (Stegomyia) aegypti* (Linnaeus, 1762), commonly referred to as the “yellow fever mosquito”, was progressively brought outside sub-Saharan Africa by human trade. It was first introduced in the Americas during the 16th century and afterwards to tropical and temperate regions of Asia and Oceania (Powell et al., 2013). Its invasion was likely favoured by a series of functional traits, such as egg desiccation-resistance, that allows them to withstand dry conditions for months, and egg moderate resistance to cold temperatures (Juliano et al., 2002; Thomas et al., 2012; Kramer et al., 2020). *Aedes aegypti* efficiently transmit several viruses to humans, including yellow fever, dengue, chikungunya, Zika, Rift Valley, Mayaro and eastern equine encephalitis viruses (Leta et al., 2018; Näslund et al., 2021; da Silva Neves et al., 2021). This is the result of several eco-evolutionary traits that are specific to the species: i) high preference for human hosts (anthropophily), which is channelled by genetic traits linked to behavioural and physiological evolutionary advantages (Harrington et al., 2001; McBride et al., 2014b), ii) exploitation of human dwellings and architectures as shelter, hide and resting indoor sites (endophily) to avoid unfavourable environmental conditions (Dzul-Manzanilla et al., 2017; Gloria-Soria et al., 2018), and iii) selection of artificial containers for oviposition and subsequent larval development (eusynantrophy; Christophers, 1960).

#### 2.1.2 Aedes albopictus

*Aedes (Stegomyia) albopictus* (Skuse, 1895), commonly referred as the “Asian tiger mosquito”, is native of tropical and subtropical regions of Southern-East Asia and Indonesia (Wat-son, 1967; Hawley, 1988). It is a competent vector of several viruses, including dengue, chikungunya, Zika, West Nile, eastern equine encephalitis and La Crosse viruses (Koch et al., 2016; McKenzie et al., 2019; Takken and van den Berg, 2019) and it was implicated as the vector species causing local transmission of dengue, chikungunya or Zika virus, even at temperate latitudes outside its native distributional range (Effler et al., 2005; Rezza et al., 2007; Delatte et al., 2008; Venturi et al., 2017; Brady and Hay, 2019; Giron et al., 2019; Barzon et al., 2021). This species is a more opportunistic feeder compared to *Ae. aegypti* (Cebrián-Camisón et al., 2020). It prefers sub-urban habitats with the presence of vegetation, dispersing bites among several species, a behaviour that might decrease the probability of pathogen transmission to humans (Turell et al., 1994; Lounibos and Kramer, 2016). Populations of this species located at temperate latitudes show: i) an adaptation to temperate climatic conditions (Marini et al., 2020) and ii) a stronger tendency to laying diapausing eggs at the end of summer (Hawley et al., 1989; Lacour et al., 2015). Diapausing eggs have been found to be resistant to below-freezing temperatures and probably allowed *Ae. albopictus* populations to overwinter and spread towards higher latitudes than *Ae. aegypti* (Hawley et al., 1989; Thomas et al., 2012).

#### 2.1.3 Aedes japonicus japonicus

*Aedes (Hulecoeteomyia) japonicus japonicus* (Theobald, 1901) [Hulecoeteomyia japonica], the “Asian bush mosquito”, originated in an area comprised between East China, East Russia and Japan (Tanaka et al., 1979a). This species may be competent for the transmission of pathogens of medical importance for humans, such as dengue, West Nile, Zika and Usutu viruses, but only experimental evidences of its role as vector exist (Takashima and Rosen, 1989; Scott, 2003; Schaffner et al., 2011; Westby et al., 2015; Veronesi et al., 2018; Jansen et al., 2018; Martinet et al., 2019; De Carlo et al., 2020; Abbo et al., 2020; Hopkins et al., 2020; but see Kilpatrick et al., 2005 for an estimated risk of transmitting WNV by this species). Its likely lesser role as a vector for human pathogens may also be assumed from the tendency to feed on other species than humans as well as the preference for more natural over urbanised areas. Established populations of *Ae. japonicus* were detected in North America from 1998 and more recently in European countries (Scott, 2003; Versteirt et al., 2009; Seidel et al., 2016; Eritja et al., 2019; Müller et al., 2020; ECDC, 2021). This species is well adapted to cold climates, overwintering either as larvae in the warmer areas, or as diapausing eggs (Krupa et al., 2021) in areas where larval habitats freeze completely (Scott, 2003; Reuss et al., 2018; Day et al., 2020).

#### 2.1.4 Aedes koreicus

*Aedes (Hulecoeteomyia) koreicus* (Edwards, 1917) [Hulecoeteomyia koreica] commonly referred to as the “Korean bush mosquito” is native to temperate areas of Northeast Asia comprising Russia, the Korean peninsula, Japan and north-east China (Tanaka et al., 1979b). This species is a suspected vector of *Dirofilaria immitis*, Japanese encephalitis and chikungunya viruses, but it has not yet been directly implicated in transmission events of zoonotic pathogens (Tanaka et al., 1979b; Montarsi et al., 2015a; Ciocchetta et al., 2018). *Aedes koreicus* is adapted to temperate climates (Versteirt et al., 2012) and has recently colonised areas of Central Europe while continuing its range expansion (Capelli et al., 2011; Versteirt et al., 2012; Montarsi et al., 2015a; Marcantonio et al., 2016; Werner et al., 2016; Negri et al., 2021; Horváth et al., 2021; Andreeva et al., 2021; Gradoni et al., 2021; ECDC, 2021). *Aedes koreicus* seems to prefer rural over highly urbanised habitats and has been found to feed on other species than humans (Montarsi et al., 2013, 2014; Cebrián-Camisón et al., 2020). In areas where *Ae. koreicus* lives in sympatry with other invasive *Aedes* species, the Korean bush mosquito is able to colonise higher altitudes and its development can start earlier in the season with respect to other AIM (Montarsi et al., 2015a; Marcantonio et al., 2016). This trait may give them a competitive advantage over other container-breeding mosquitoes whose adults emerge later in the season.

### 2.2 The theoretical structure of the model

The basic structure of `dynamAedes` has been described in Da Re et al. (2021). We amended some components of the model to generalise its structure. Thus, we provide here a short recap of model structure while describing the new model features.

`dynamAedes` is composed of three main compartments (life stages) that represent a simplified version of *Aedes* the mosquito life cycle: egg, juvenile and adult stages (Fig. 1). Larval and pupal stages, which can be assumed to have somewhat similar physiological requirements, are fused in a unique “juvenile” compartment. Each compartment is divided into sub-compartments to account for the different physiological states for individuals in the three main compartments (e.g. 1-day old adult females that are not sexually mature). The number of sub-compartments into each compartment is dictated by the known minimum number of days needed by each species to pass to the next stage or complete the gonotrophic cycle (for adults). Thus, the minimum duration of development in each compartment varies among developmental stages as well as among species. As an example, the whole duration of the developmental cycle (i.e. from egg-laying to adult emergence) has a minimum duration of 11 days for *Ae. aegypti* and *Ae. albopictus*, whereas 21 days for *Ae. koreicus* and *Ae. japonicus* (see Tab S3 in SM for generic model assumptions).

**Figure 1:**
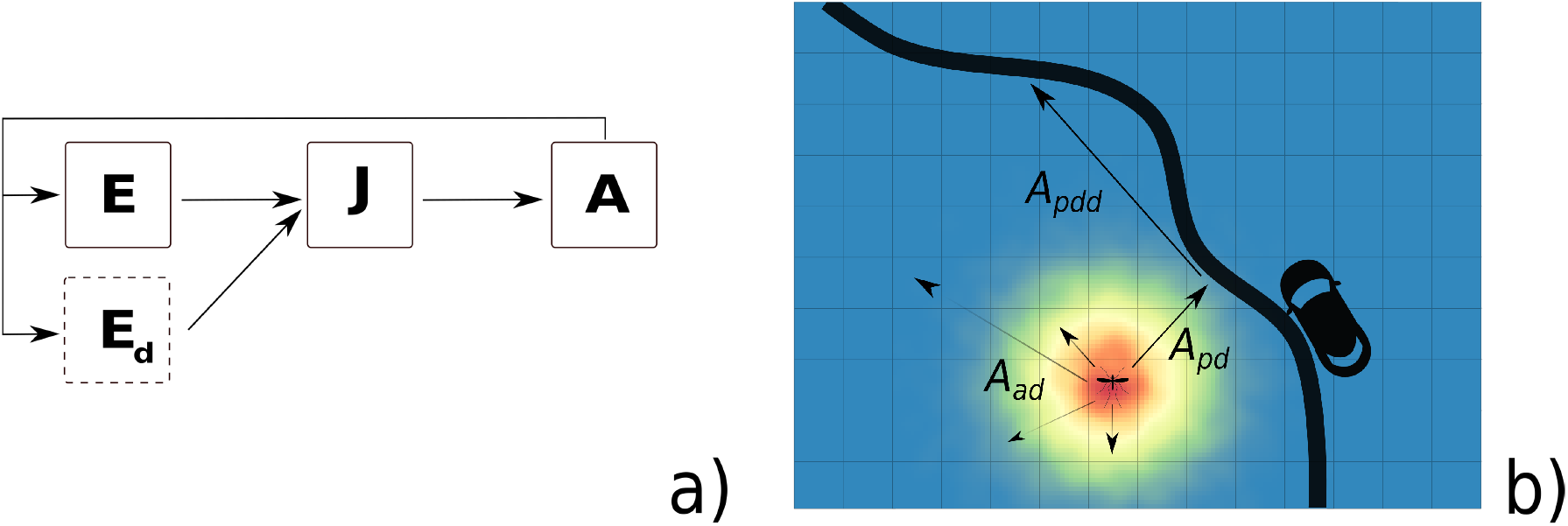
Graphical representation of `dynamAedes` model structure, adapted from Da Re et al. (2021): a) the life cycle of a generic simulated mosquito species, while in b) a representation of active and passive dispersal processes happening within the Adult (A) compartment at local scale. E: egg compartment; Ed: diapause egg compartment (available for all species except *Ae. aegypti*); J: juvenile compartment; A: adult compartment; Aad: adult active dispersal; Apdd: adult passive dispersal; Apd: adult probability of being caught in a car.

In the model, time is treated as a discrete quantity and “day” is the fundamental temporal unit. Therefore, each event in the simulated life cycle occurs once per day and always in the same order. The model can be run with or without a spatial structure. If the model is spatially explicit, space is treated as a discrete quantity. In this case, the fundamental spatial unit is a (user defined) cell of a lattice grid into which the species life cycle takes place and, if relevant (see below), among which adult mosquitoes disperse.

Adult female mosquitoes lay non-diapausing eggs, E, in the summer months or diapausing eggs, Ed, at the end of the season. The embryonic development and hatching of diapausing eggs are activated by increasing daily temperature and photoperiod (typically at the end of winter or early spring). All the developmental and reproductive events considered in the model were treated as stochastic processes with probabilities derived from temperature(or photoperiod)-dependent functions by following the generic formulation:

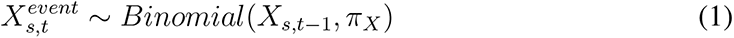

where *X_s,t−1_* may represent eggs, juveniles or adults that undergo one of the following events in the life cycle: lay eggs, hatch, emerge or survive in cell *s*, at the end of the day *t –* 1. *π_X_* is the temperature-dependent (or photoperiod dependent for the hatching of diapausing eggs) daily probability of any of the life cycle events *X*. All the temperature-dependent functions were calibrated using data from the scientific literature (see Tab S4 in SM) fitted using exponential, polynomial equations, and non-linear Beta density functions, using a combination of `drc` (non-linear models) and `aomisc` (Beta function self-starters) R packages (Ritz et al., 2015; Onofri, 2020). The beta function derives from the beta density function and it has been adapted to describe phenomena taking place only within a minimum and a maximum threshold value (threshold model), such as physiological rates with respect to temperatures in the mosquito life cycle (Onofri, 2020). In a similar fashion, adult active dispersal was modelled as species-specific log-normal decaying functions of distances derived from dispersal estimates from field observations for *Ae. aegypti* and *Ae. albopictus* (Roche et al. 2015; Marcantonio et al. 2019; Marini et al. 2019a; Müller et al. 2020; see Tab S5 in SM for dispersal parameters). In addition to active dispersal, the model also considers dispersal aided by cars along the main road network (a matrix containing the coordinates of the grid cells of the landscape intersecting the road network must be provided, see the “Spatial scales of the model and temperature data sources” section), defined as the “hitchhiking” probability of a female to enter in a car and to be driven and released further away. This probability has been defined for all species by estimates measured for *Ae. albopictus* (Eritja et al., 2017), while the average distance covered by a single car trip was taken from Pasaoglu et al. (2012). This type of dispersal is thought to be amongst the main drivers of medium-range geographical expansion for invasive *Aedes* mosquitoes, especially for *Ae. aegypti* and *Ae. albopictus* (Marcantonio et al., 2016; Eritja et al., 2017; Müller et al., 2020).

Density-dependent survival is an important regulatory factor of mosquito population dynamics (Gilpin and McClelland, 1979). Its regulatory effect for juvenile stages appears to be more common in mosquitoes breeding in container or highly ephemeral habitats (Juliano, 2007), such as invasive *Aedes*. In `dynamAedes` we parameterised a density-dependent function by extracting observations on *Ae. aegypti* from Figures 2a and 2b in Hancock et al. (2016) using the Webplot Digitizer (Rohatgi, 2020). We considered the proportion of juveniles that survived through the juvenile stage (in a 2 L container) reported by these authors as an estimate of juvenile survival probability at different densities. Mortality probabilities (1 – *proportion surviving*) were converted into rates, which were scaled to a daily time step dividing by the corresponding immature development time (in days) at different densities. Finally, we regressed the natural logarithms of these daily mortality rates on the corresponding densities. The fitted daily survival rate at different densities was then summed to the temperature-dependent juvenile mortality. The resulting probability was then used to inform a binomial random draw (see equation 1) describing overall juvenile daily survival.

**Figure 2:**
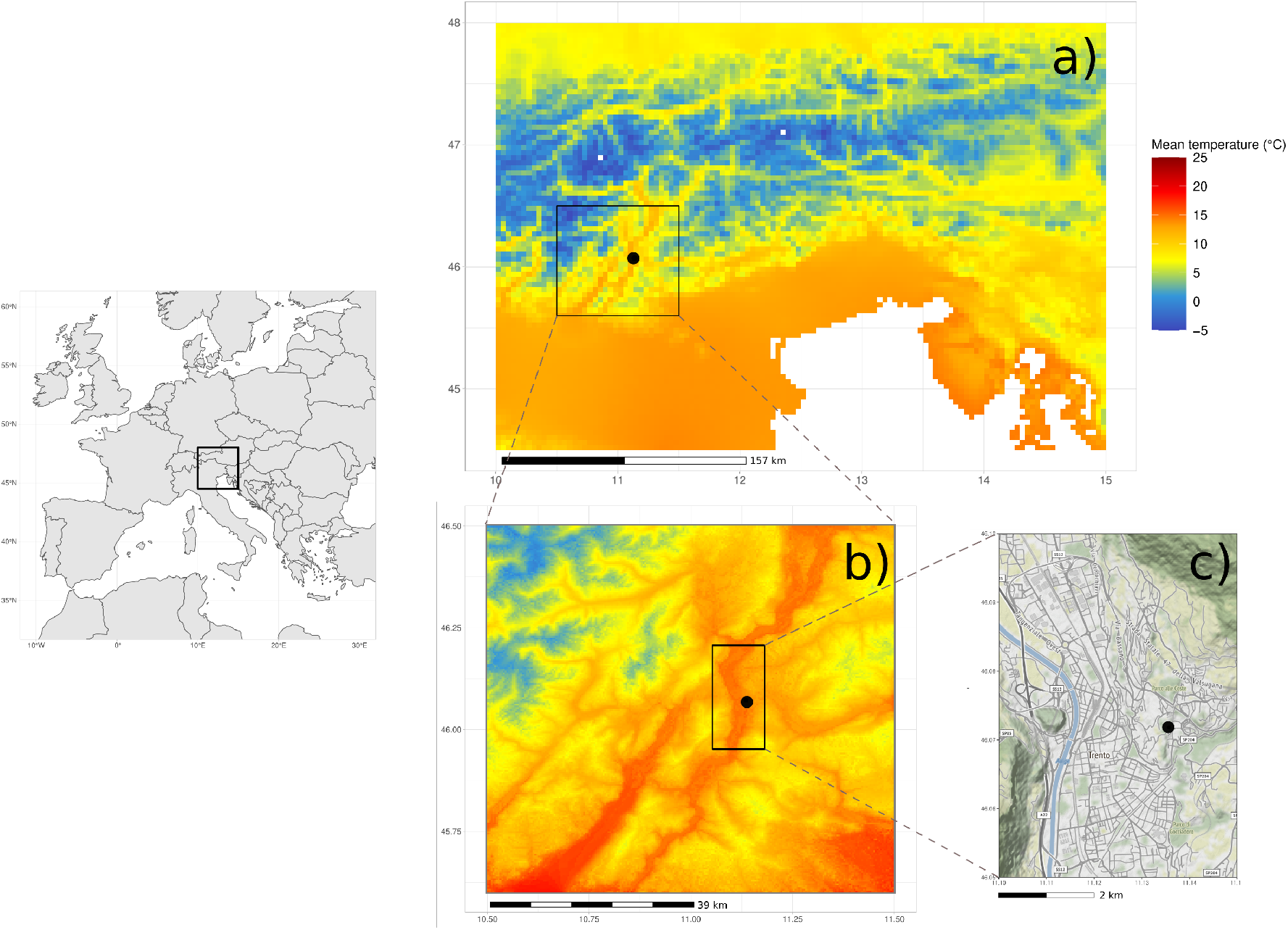
`dynamAedes` allows to simulate *Aedes* mosquitoes population dynamics at three different spatial scales: a) regional, b) local, and c) punctual (weather station). Passive and active dispersal is enabled only at local spatial scale.

Some invasive *Aedes* can lay eggs resistant to low temperature commonly referred to as “diapausing eggs” (Thomas et al., 2012; Lacour et al., 2015; Krupa et al., 2021). Diapause describes the evolutionary adaptation exploited by insect species to overcome unfavourable environmental conditions by passing through an alternative and dormant physiological stage. In *Ae. albopictus*, maternal photoperiod is the environmental stimulus implied to induce oviposition of “diapausing eggs” (Pumpuni et al., 1992; Lacour et al., 2015). In `dynamAedes`, the oviposition of diapausing/non-diapausing eggs was integrated as a species-specific exponential function on the incidence of diapausing eggs given photoperiod length (and thus geographically-dependent; Urbanski et al. 2012; Petrić et al. 2021). The function is based on data from Lacour et al. (2015) for *Ae. albopictus* and Krupa et al. (2021) for *Ae. japonicus*. We applied the same diapausing function developed for *Ae. japoncus* to *Ae. koreicus* due to the close phylogeny of these species and the lack of data for *Ae. koreicus* (see Tab. S6). The daily survival of diapausing eggs was set to be constant (0.99) only for *Ae. japonicus* and *Ae. koreicus*, while for *Ae. albopictus* we used the exponential function described in Metelmann et al. (2019). The hatching rate of diapause eggs was triggered by an increasing photoperiod regime (spring) from 11.44 hours of light for *Ae. albopictus* (95th percentile estimated from Petrić et al. 2021) and 10.71 hours for *Ae. japonicus* or *Ae. koreicus* (Krupa et al., 2021).

### 2.3 Overview of the R package

The function `dynamAedes.m` calls the model and allows to customise the simulated scenario through a suite of options. As for the simplest application of the model (no explicit spatial dimension, scale=“ws”, see next paragraph for further details), the user has to define what species to model through the argument species (default “aegypti”), the “type” and number of introduced propagules through intro.eggs, intro.juvenile or `intro.adults` (default `intro.eggs`=100, `intro.juvenile`=0, `intro.adults`=0), and the volume (L) of water habitats wanted in each spatial unit with the argument `lhwv` (larval-habitat water volume, parameterised from Hancock et al. 2016; default `lhwv`=2; see Fig S13 for a sensitivity analysis of this parameter). Moreover, the argument `temps.matrix` takes the matrix of daily average temperature (in Celsius degree) used to fit the life cycle rates. This matrix must be organised with the daily temperature observations as columns and the geographic position of the *i*-grid cell as rows (it follows that the matrix will have only one row when `scale=“ws”`). The day of start, end and number of iterations are defined by `startd`, `endd` and `iter`, respectively. The model has been optimised for parallel computing and the number of parallel processes can be specified through the option `n.clusters`. If the modelled species is *Ae. albopictus*, *Ae. japonicus* or *Ae. koreicus* (e.g., `species=“albopictus”`) then the arguments defining latitude (`lat`), longitude (`long`) and year of introduction (`intro.year`) should be adequately defined to allow a correct switch to and from the egg diapausing stage.

The default output of dynamAedes consists of a list of numerical matrices containing, for each iteration, the number of individuals in each life stage per day (and for each grid cell of the study area if `scale=“lc”` or `“rg”`). If the argument `compressed.output=FALSE` (default TRUE), the model returns the daily number of individuals in each life stage sub-compartment. The model, coded in the R statistical language (R Core Team, 2021), and adapted for parallel computation, is available on the the following link https://github.com/mattmar/dynamAedes (it has meanwhile been submitted to the CRAN).

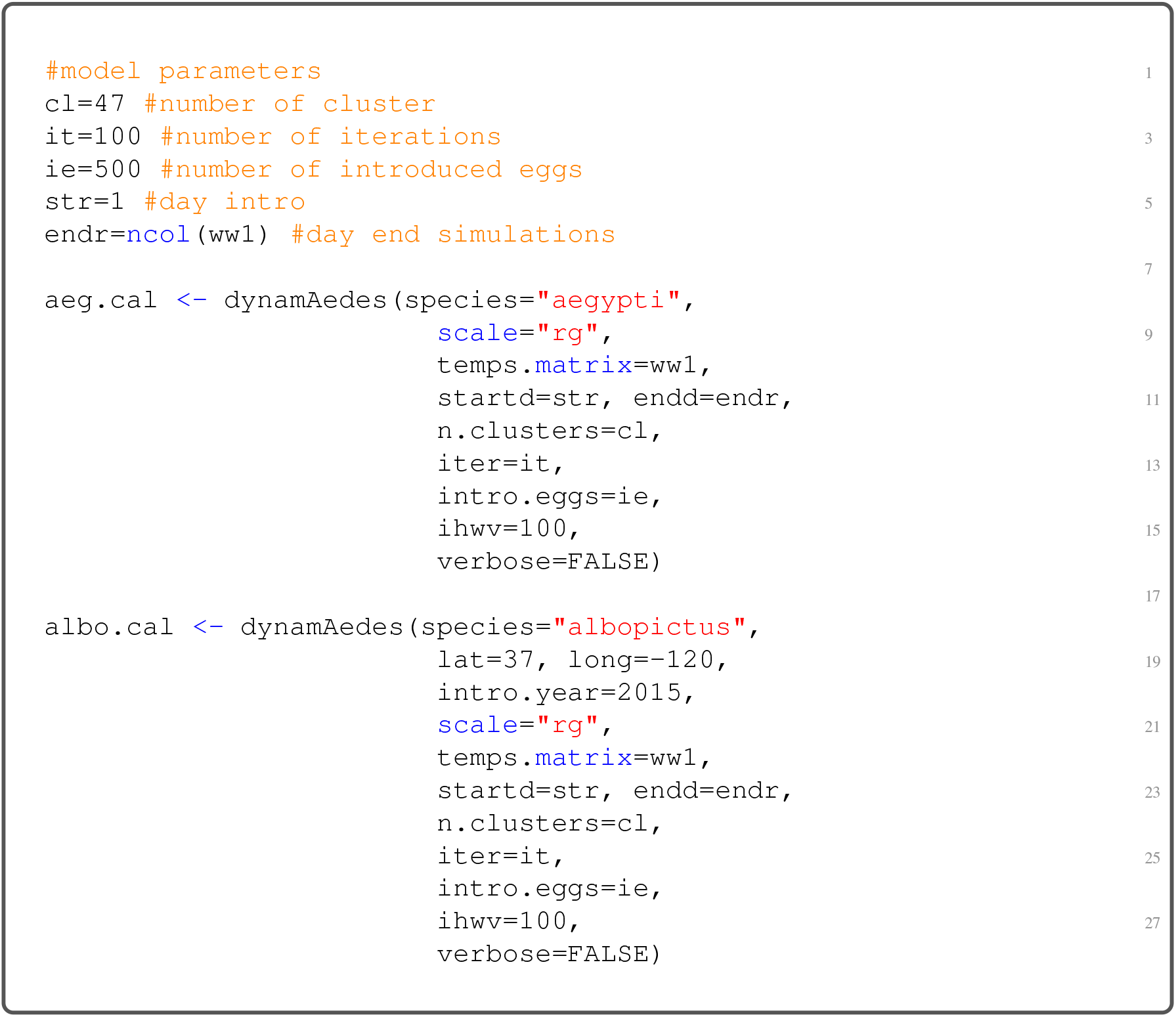

#### 2.3.1 Spatial scales of the model and input temperature data

The selection of the geographical scale for population dynamics is a crucial aspect of the whole package and the temperature dataset provided to `dynamAedes` function must reflect this decision. Along with the photoperiod, temperature is the other only environmental driver of our model, which is dictated by its central role in mosquito development and activity. The measurement of temperature is inevitably scale-dependent, thus we structured the model to allow for temperature datasets relevant for different measurement spatial scales (Fig. 2) and to match the different hypotheses that users may want to test.

The punctual or “weather station” scale (`scale=“ws”`) is the smallest geographic scale (i.g., no spatial dimension) available in `dynamAedes` and the environment modelled is assumed to be what represented by the chosen weather station (or any other data-loggers). In this case, the model has no spatial structure, thus dispersal is not considered: the model will return the temporal trend of population dynamics given the chosen temperature, larval-habitat water volume and photoperiod conditions.

The “local” scale (`scale= “lc”`) represents those scenarios and spatial resolutions at which species dispersal and local microclimate variability are relevant for users. We suggest to keep the resolution of the matrix of temperatures equal or smaller than the maximum daily dispersal range of the mosquito species (i.e., usually under 1 km for *Aedes* species; Guerra et al., 2014). The optional arguments `cellsize`, `dispbins` and `maxadisp` are available to fine tune the dispersal kernel which drives the spatial behaviour of the simulated mosquito populations. The argument `cellsize` (default `“cellsize=250”` meters) sets the minimal distance of the dispersal kernel and should match the size of the cell to avoid inconsistencies (i.e., mosquitoes dispersing at a finer or bigger grain than the arena), `maxadisp` sets the maximum daily dispersal (default `maxadisp=600 meters`), and `dispbins` the resolution of the dispersal kernel (default `dispbins=10`). Passive dispersal is also implemented and it requires i) a matrix containing the coordinates of the grid cells of the landscape intersecting the road network (argument `road.dist.matrix`), and ii) to specify the average car trip distance through the argument `country`, which can be defined by the user or considering estimates for the following countries: France, Germany, Italy, Poland, Spain and the United Kingdom Pasaoglu et al. (2012). An extensive example of model application at local spatial scales is described in Da Re et al. (2021).

The rationale behind the third spatial scale considered in the model, the “regional” scale (`scale= “rg”`) was to return an overview of invasive *Aedes* population dynamics over large extents (i.e., larger than 1 km). The model in regional scale does not account for species dispersal, introductions happen separately (but at the same time) in each grid cell which hence are closed systems. The output of the model at “regional” scale can be compared to those produced by correlative species distribution models (SDMs), with the advantage of mechanistic rather than purely correlative model foundations.

The amount of water available for larval development in each spatial unit(s) (at any model spatial scale) was set as 2 L that is the water volume considered in the experiments we used to parametrise model functions Hancock et al. (2016). It is likely that for many real-world model applications, the relative availability of breeding habitats is much higher, and we encourage users to set a value based on their scenarios and hypotheses (i.e. through the model option `lhwv`; Hartemink et al. 2015).

#### 2.3.2 Auxiliary functions

Several auxiliary functions are available to analyse model outputs. The function `psi` returns the proportion of model iterations that resulted in a viable population for the given date. It works for all spatial scales and the output can reflect either the overall grid or each single cell. Likewise, summaries of mosquito abundance at each life stage for each day can be obtained through `adci`, which by default returns the inter-quartile range abundance of each life stage. Similarly, `icci` returns a summary of the number of invaded cells over model iterations. Estimates of dispersal spread (in km^2^) of the simulated mosquito populations is provided by the function `dici`, which is available only for model results computed at the local scale (the only scale which integrates dispersal). Finally, the function `get_rates_spatial`, returns the output of the temperature-dependent physiological functions used by `dynamAedes` to derive the daily rates. It can be used to better understand the outcome of model simulations, by highlighting those areas where the predicted values of the temperature-dependent functions are maximised or minimised, or to derive causal-based physiological estimations that, for example, could be used as inputs for correlative SDMs (e.g., Kearney and Porter, 2009; Mathewson et al., 2017).

## 3 Case studies and model validation

We applied the model to three case studies representing different geographical scales and areas, species and invasion trajectories. The case studies were chosen considering the availability of optimal mosquito data-sets to show model strength and weakness. We did not report any example for the “local scale” as it had already been provided in Da Re et al. (2021), who applied a previous version of `dynamAedes`.

### 3.1 Regional scale models

#### 3.1.1 Likelihood of successful introductions of *Ae. albopictus* and *Ae. aegypti* across California, USA

We used `dynamAedes` at “regional” scale to assess the likelihood of successful introductions for two invasive *Aedes* across California: *Ae. aegypti* and *Ae. albopictus* which are considered established from 2011 and 2013, respectively (Fujioka, 2012; Gloria-Soria et al., 2014). California is among the few areas where established populations of these two species were detected during the last decade and their progressive spread was documented in great detail^2^. We downloaded daily minimum and maximum temperature data from the NASA Daily Surface Weather Data on a 1-km Grid for North America (DAYMET), Version 4 (Thornton et al., 2020) from 1 January 2011 to the 31 December 2018. These two sets of data (in netCDF raster format) were clipped to the boundary of California and aggregated at a spatial resolution of 2.5 km by using a combination of GDAL (GDAL/OGR contributors, 2021) and Climate Data Operators (CDO; Schulzweida, 2019) software. The two sets of raster layers were then imported in R 4.0.4 (R Core Team, 2021), transformed in matrices, averaged and converted to integers to obtain a single average daily temperature integer matrix with cell id as observations (rows) and days as variables (columns). This dataset was used as the input temperature matrix for the model. We run 80 model iterations for five years, introducing 500 eggs in each cell of the gridded landscape on 15 May 2011 and 2013 for *Ae. aegypti* and *Ae. albopictus* respectively. By using the auxiliary function `psi`, we then derived a map showing the proportion of iterations with viable population of both *Ae. aegypti* and *Ae. albopictus* at the end of the simulated period (15 May 2016 and 2018). The photoperiod was set to match conditions in the geographical center of California (Lat 37° N, Lo -120° W) and the amount of breeding habitat was set to 100 L (per cell), which we assumed to be representative of the potential water larval habitats available given cell size and overall regional climate.

We validated model prediction using maps of cities in California with known species presence updated to 2021 by the California Department of Public Health (CDPH^3^). We derived the average predicted probability of successful introduction for each city and computed the Area Under the ROC Curve (AUC) score which defines the probability that a randomly chosen positive city will be ranked higher than a randomly chosen negative city. An AUC score > 0.5 indicates that the model is performing better than random, while a score of 1 indicates perfect prediction. In addition, we calculated the percentage of positive cities that fell into a grid cell that had a probability of establishment higher or equal to 1% (e.g., at least 1 out of the *n_th_* iterations reported a viable mosquito population in that cell).

The predicted spatial pattern of areas with a high likelihood of *Ae. aegypti* and *Ae. albopictus* successful introduction is in general consistent with updated information on the presence of this species (Fig. 3). Predictions show moderate-to-high probabilities of a successful introduction in all counties with known presence of these species, except for the extreme South-East coastal part of the state, where *Ae. aegypti* was predicted to have a low probability of successful introduction whereas being well established (Fig. 3). This may be due to the high micro-climatic variability that characterises coastal areas of California which may not be resolved by the temperature datasets that we have considered in this case study. On the contrary, areas predicted to be suitable for *Ae. albopictus* exceeded by far the known current distribution of this species. Factors other than temperature and photoperiod, more nuanced aspects of species invasion history and the extremely low humidity during the dry season in the Central Valley of California or the Inland Empire, may hinder species establishment in these areas. Nevertheless, recently *Ae. albopictus* was found as north as Redding, Shasta County, thus it is not unlikely that this species is also present (perhaps at low densities), but not yet detected, at southern latitudes in California.

**Figure 3:**
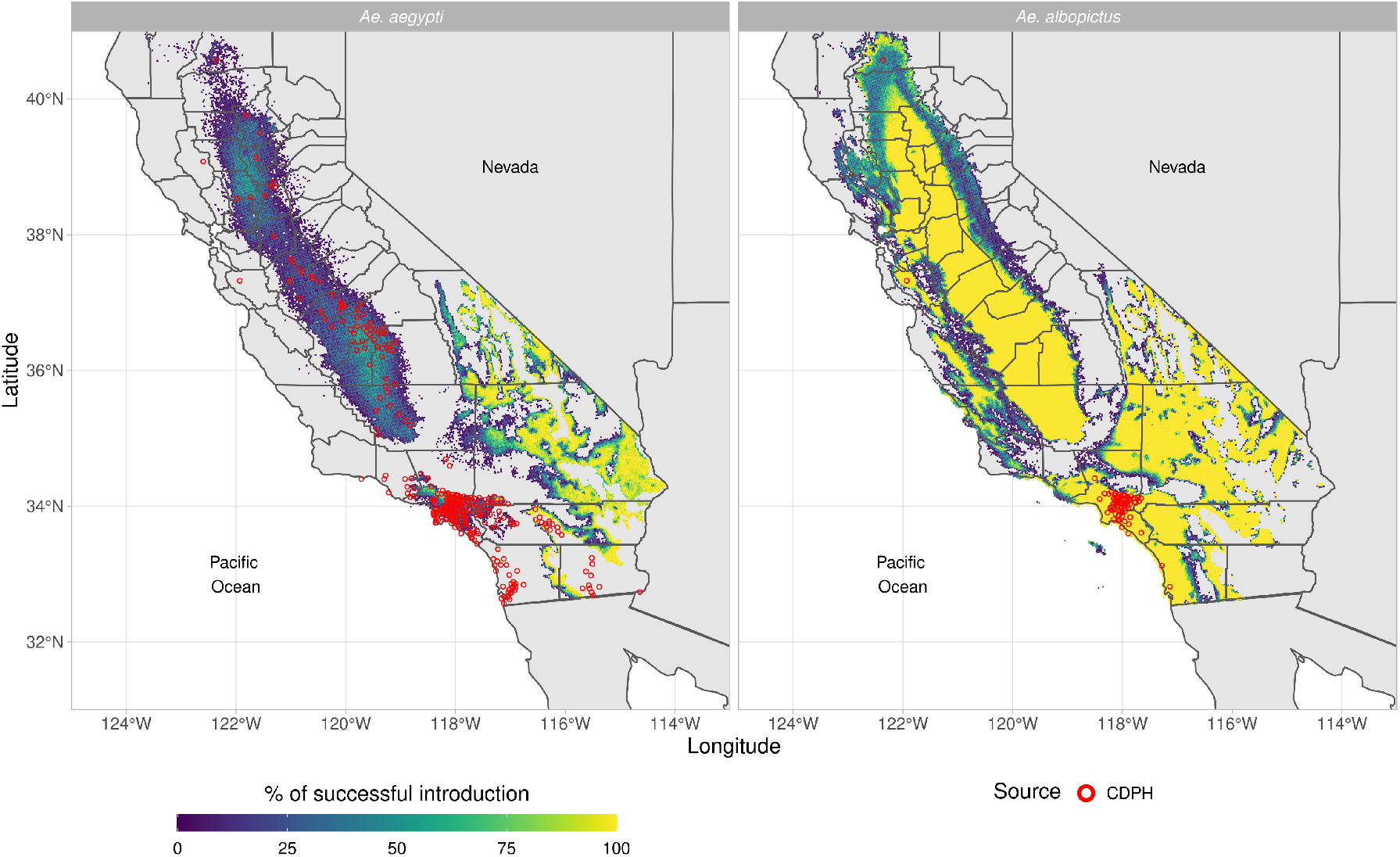
Predicted percentage of established introductions of *Ae. aegypti Ae. albopictus* in California (USA) for the years 2011-2016 and 2013-2018, respectively. The red dots represent the centroids of the Californian municipalities with established populations as reported by the Californian Department of Public Health (CDPH).

Both models had over 75% of successful introduction scores (calculated as the proportion of pixels with species observations and simulated proportion of invasion > 1%) when validated at city levels (Tab. 1). Still, only the prediction for *Ae. aegypti* validated at city levels reported an AUC score bigger than 0.5 (Tab. 1), whereas validating the model by averaging the predictions at county level resulted in an AUC bigger than 0.5 for both species (0.892 and 0.717 for *Ae. aegypti* and *Ae. albopictus* respectively; Fig. S14).

**Table 1:** Model validation. The column “Sensitivity 1%” reports the proportion of cities predicted to have at least one successful introduction over the total number of iterations (predicted introduction success equal or over 1%. *** p.value < 0.001).

| Species | Geographic location | Scale | AUC | Specificity | Sensitivity | Sensitivity 1% | Spearman's $\rho$ |
| --- | --- | --- | --- | --- | --- | --- | --- |
| *Ae. aegypti* | California | Regional | 0.658 (0.401-0.914) | 0.8 | 0.642 | 0.795 | - |
| *Ae. albopictus* | California | Regional | 0.435 (0.367-0.502) | 0.241 | 0.641 | 1 | - |
| *Ae. albopictus* | France | Regional | 0.874 (0.867-0.880) | 0.788 | 0.802 | 0.848 | - |
| *Ae. albopictus* | S-E France | Punctual | - | - | - | - | 0.753*** |
| *Ae. koreicus* | N-E Italy | Punctual | - | - | - | - | 0.735*** |

#### 3.1.2 Likelihood of successful *Aedes albopictus* introductions in France

*Aedes albopictus* was first detected in metropolitan France in 1999 (Schaffner et al., 2000) and since 2004 it has established populations in the southern part of the country while still expanding its distribution range (ECDC, 2021). We used `dynamAedes` model at “regional” scale to assess the success of introductions of *Ae. albopictus* for the whole metropolitan France. We processed ERA5-Land (Muñoz-Sabater et al., 2021) hourly air temperature measured at 2 m above surface from January 1st 2015 to December 31st 2020 in the Climate Data Store (CDS) Toolbox^4^ to get the daily mean temperature of the period considered for the whole metropolitan France, at the spatial resolution of ∼ 9 km. The netCDF file obtained was imported in R 4.0.4 (R Core Team, 2021), where was clipped to the extent of metropolitan France, converted from degrees Kelvin to Celsius and converted to integer to obtain a single average daily temperature integer matrix with cell id as observations (rows) and days as variables (columns). This dataset was then used as the input temperature matrix for the model. We run 100 simulations for five years, introducing 100 eggs in each cell of the gridded landscape on 15 May 2015. By using the auxiliary function `psi`, we derived a probability map of the areas showing the proportions of iterations that produced a viable population of *Ae. albopictus* five years after the simulated introductions.

We validate the model predictions using the list of the 3419 French municipality reporting established *Ae. albopictus* until 2020 provided by the French Health Ministry (data collected thanks to active monitoring from French mosquito operators and passive surveillance^5^). Similarly to the previous case study, we computed both the AUC as well as the proportion of positive location falling into a grid cell that had a percentage of established introductions higher than or equal to 1%. The spatial pattern of the areas predicted to have a high likelihood of successful *Ae. albopictus* introduction (Fig. 4) is consistent with updated observational data (*Ae. albopictus* map; ECDC, 2020) as well as with the results of other mechanistic models (see for instance Metelmann et al., 2019; Pasquali et al., 2020). The Mediterranean French coast and the Rhone valley are the areas where our model predicted the highest percentage of successful introduction. Similarly, the Aquitaine region on the Atlantic coast and the Alsace region in the North-East part of France showed relatively high predicted percentage of successful introduction. The northern and the central part of France, as well as the Pyrenees areas, show low percentage of successful introduction under the current climatic conditions. However, the resolution of the pixel, approximately 10 km, may have played a role influencing the model outcomes especially in topographically complex areas such as the Pyrenees or the French Alps, where the microclimate of the valleys may be underestimated. Similarly, the model was not able to predict the successful introduction of the species in areas such as Paris, where *Ae. albopictus* is established and probably favoured by i) local climatic factors such as the urban heat island effect, and ii) the continuous inflow of *Ae. albopictus* propagules to Paris from areas where the species is already established. Indeed, most railways, flights and highways have a connection with Paris, and the Paris-Lyon-Mediterranée axis is the main artery of France, with an average of >60,000 cars/day on the highway^6^ and 240 trains per day^7^, thus the quantity of imported mosquitoes can compensates for the less favorable climatic conditions of Paris compared to the Mediterranean region (recent phylogeographic findings support this hypothesis, see Sherpa et al., 2019). All the model performances metrics assessed support the capacity of the model to discriminate between areas where the species can or cannot be established (Tab. 1).

**Figure 4:**
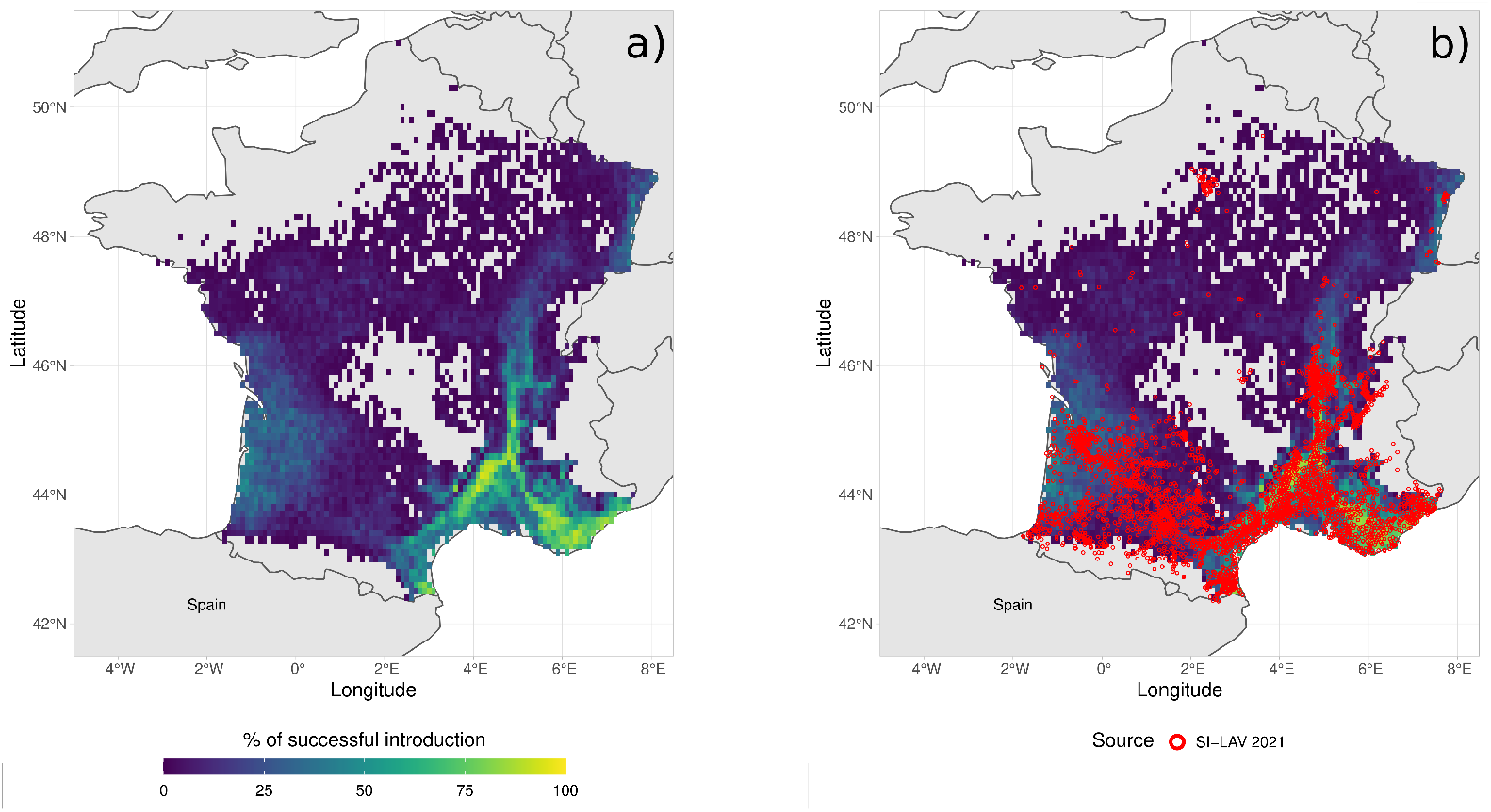
Percentage of successful introduction for *Ae. albopictus* in France for the years 2015-2020: a) Model prediction, b) Model prediction and in red the centroids of the French Municipalities with established population of *Ae. albopictus* reported by the French Health Ministry (SI-LAV).

### 3.2 Punctual population dynamics temporal trends

#### 3.2.1 *Aedes albopictus* population dynamics in Nice, South-East France

*Aedes albopictus* is established mainly in the southern part of metropolitan France where more than 40% of municipalities are colonized^8^ and it is still expanding in other areas. We computed the population dynamics of the species informing `dynamAedes` model at punctual spatial scale using temperature observations downloaded from the National Oceanic and Atmospheric Administration (NOAA) network via the R package `rnooa`. The observations of the NOAA weather station located in Nice area (usaf code = 076900, Lat 43° 42’ 00” N, Lon 7° 13’ 12” E, 3.7 m a.s.l.), spanning from January 1st 2013 to December 31st 2018, were linearly interpolated to fill missing values at hourly and daily level. Afterwards, the daily average for all observations was computed. We run 100 simulations for five years, introducing 500 eggs on 15 May 2013. By using the auxiliary function `adci`, we then derived the daily abundance inter-quantile range for each life stage and the abundance of newly-laid eggs per day.

The model was validated comparing the simulated newly laid eggs per day to egg counts from mosquito ovitrap data, following the validation approach presented in Tran et al. (2013). During the years 2014-2018, fifty ovitraps were installed in the Nice area and inspected fortnightly from April-May to November-December (data collected and kindly provided by EID Méditerranée). We computed the Spearman’s *rho* correlation coefficient between the weekly-aggregated simulated newly-laid eggs and the mean observed eggs per day.

Results showed that the model was able to correctly reproduce the seasonal dynamics of the new-laid eggs over five years (Spearman’s *rho* = 0.753, p.value < 0.001). The first simulated eggs were laid during the late spring each year, confirming the fact that the first overwintering eggs hatch at the end of the winter season or at the beginning of spring. The ovipositing season seem to last until the late autumn accordingly to the observations, while our model seems to predict a shorter length of the ovipositing period. Nevertheless the model is able to correctly infer the peak of the ovipositing season during the summer months (Fig. 5).

**Figure 5:**
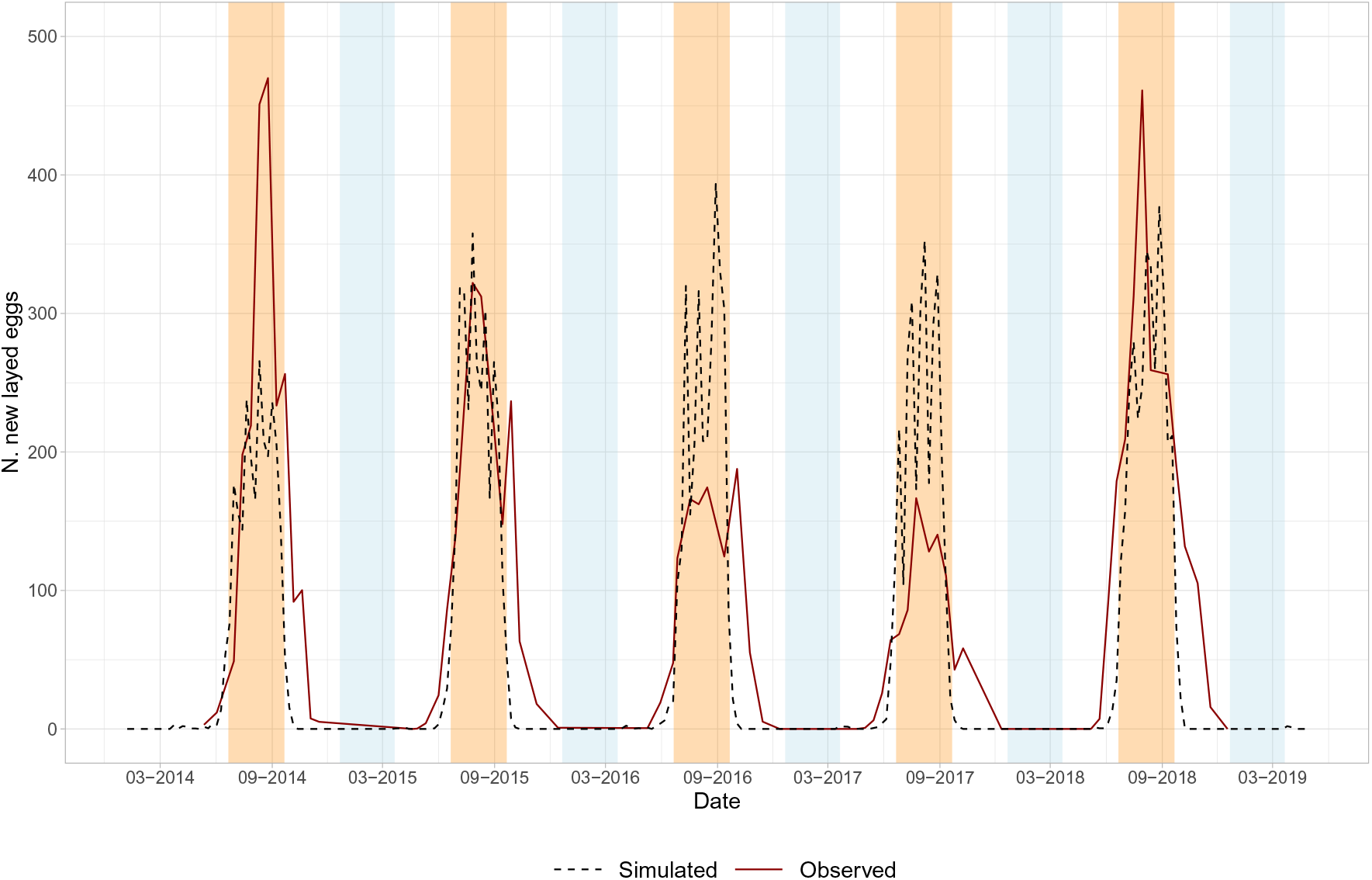
Temporal trend reporting the simulated and observed new-laid eggs of *Ae. albopictus* years 2014-2018 in Nice, SE France. The light blue bands represent the winter seasons, while the orange bands the summer seasons. The simulated data were rescaled for graphical purpose using as rescaling factor the ratio between the maximum observed value and the maximum median simulated values.

**Figure 6:**
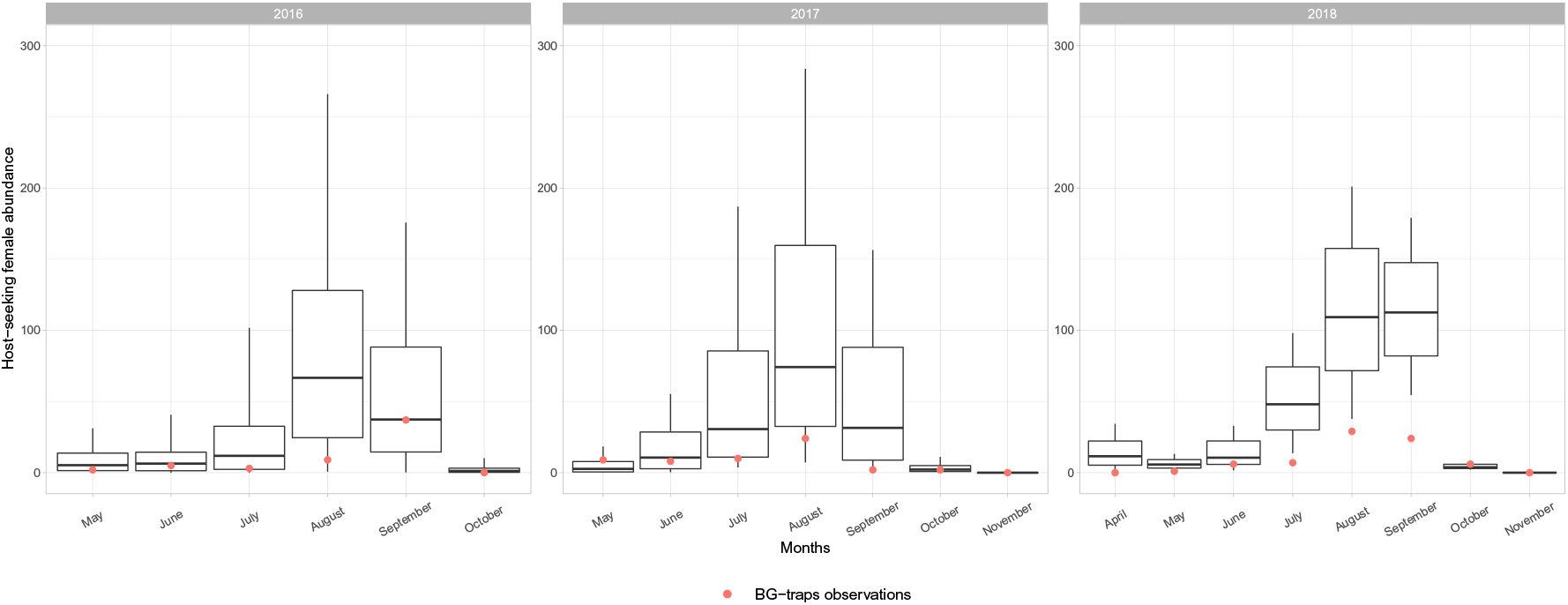
Temporal trend reporting the boxplot of simulated and observed host-seeking *Ae. koreicus* females for the years 2016-2018 in Trento, NE Italy.

#### 3.2.2 *Aedes koreicus* population dynamics in Trento, North-East Italy

*Aedes koreicus* was first detected in Trento Autonomous Province (NE Italy) in 2013, soon after the first Italian detection in the neighboring Belluno province (Capelli et al., 2011).

We computed the population dynamics of the species informing `dynamAedes` model at punctual spatial scale using temperature observations downloaded from the local network of weather stations^9^. The daily average temperature observations from the “Trento Laste” weather station (Lat 46° 04’ 18.5” N, Lon 11° 08’ 08.5” E, 312 m a.s.l.) spanning from 1 January 2015 to 31 December 2018 were linearly interpolated to fill missing values. We run 100 simulations for five years, introducing 500 eggs on 15 May 2015. Using the auxiliary function `adci`, we then derived the daily inter-quartile range abundance for each life stage and for the daily host-seeking female sub-compartment.

The model was validated computing Spearman’s *rho* correlation coefficient between the monthly-aggregated simulated host-seeking females and the observations gathered from four BG-Sentinel traps installed in Trento municipality from April to November during the years 2016, 2017, and 2018 (data obtained from Marini et al. 2019b). In order to compare observed and simulated data, the whole simulated host-seeking females abundance was multiplied by a BG-sentinel catching rate equal to 0.157, as estimated by Marini et al. (2019b) and similar to what already reported for *Ae. albopictus* in previous studies (Guzzetta et al., 2017).

The simulated population dynamics showed that *Ae. koreicus* could be successfully introduced in the study area. The model correctly predicted the start of the seasonal activity in early spring, while the higher abundance of female host-seeking mosquitoes was predicted to be in late summer. Considering the three years together, our model was able to reproduce the observed seasonal population dynamics, where the 76.2% (47.6%) of the observed captures lie within the 95% (50%) credible intervals of model predictions (Tab S7). Similarly, the Spearman’s rho for the three yeas was 0.735 (p.value < 0.001) (Tab. 1)

## 4 Discussion

From an ecological perspective, our modelling approach focuses on the species fundamental thermal niche (*sensu* Hutchinson, 1957), since we considered temperature as the main driver of population growth and dynamics. In light of this, our model was able to infer spatial and temporal population dynamics of different species, at different geographic scales and locations. The model showed overall good validation performance, and the areas predicted to likely support *Aedes* mosquito populations largely matched what reported by observational studies and other existing models (Kraemer et al., 2015, 2019; Oliveira et al., 2021).

Nevertheless, we raise the attention on two aspects that should be considered when applying `dynamAedes` and interpreting model results. First, good quality information on survival and developmental rates is largely available for *Ae. aegypti* and *Ae. albopictus*, whereas much less for the other two species. Similarly, mosquito observational datasets with sufficient longitudinal depth for model validation are scarce for *Ae. koreicus* and presently absent, to the best of our knowledge, for *Ae. japonicus*. Thus, whilst we have built the foundations for an open-source modelling framework that can be progressively expanded, life cycle functions and thus outputs for *Ae. japonicus* and *Ae. koreicus* should be interpreted more carefully.

Secondly, we recognise that pixel size may influence model outcome because of the aggregating effect of the Modifiable Unit Areal Problem (Jelinski and Wu, 1996; Da Re et al., 2020). While the consequences of this artifact are well known in SDMs applications, they are rarely mentioned or addressed (but see Peterson, 2014). Thus, the correct choice of temperature datasets is crucial to investigate species population dynamics and interpret model results (Bütikofer et al., 2020). Climatic reanalysis and Global/Regional Circulation Models are reliable data sources with high temporal resolution for present climatic conditions and robust future projections, though they have a coarse spatial resolution that may underestimate the effect of microclimate on species biology (Metelmann et al., 2019; Liu-Helmersson et al., 2019). Coarse resolution of input temperature data is rather common since temperature is often estimated over large extents, and can become an issue in topographically complex regions where the effect of microclimatic variation on population dynamics may pass undetected. The results showed in Fig. 3–4 may be better interpreted considering the coarse resolution of temperature data which may have caused lower proportion of established populations in topographically complex areas such as the Pyrenees or the Peninsular Ranges. Interpolated local micro-climatic conditions, for example estimated with the `microclima` R package (Maclean et al., 2019), have the advantage of providing fine spatial and temporal resolution datasets. But, they need to be first properly validated with field data which are typically difficult to gather also over small geographical extents. Temperature measured with classical weather stations may be considered as the most accurate available observations of local climatic conditions. Though it is not always easy to deal with such data due to the limited number of weather stations and gaps in the time series, they are suited for statistical downscaling or bias adjustment for climate change projections (Bedia et al., 2020).

### 4.1 Model assumptions

Model structure has been designed to be as ecologically relevant as possible considering available data, however, when data were limited, we relied on a set of “expert-based” assumptions that must be clearly stated.

The interplay of multiple environmental factors drives the population dynamics of *Aedes* mosquitoes but we chose to mold our model framework just on established information available for temperature and photo-period (Pumpuni et al., 1992; Waldock et al., 2013; Eisen et al., 2014). This choice was suggested by a generalised lack of clear relationships between other environmental factors and *Aedes* population dynamics. For example, concerning the role of precipitations, different studies report contrasting results (Koenraadt and Harrington, 2008; Tran et al., 2013; Caldwell et al., 2021). Moreover, invasive *Aedes* mosquitoes mostly thrive in urban or suburban landscapes where the presence of standing water is often independent from precipitations (except for extreme rainfall events (Roiz et al., 2015). We suggest that, at the present stage, `dynamAedes` it is better suited for applications in temperate climates, where temperature seasonality is a more important limiting factor than in tropical climates, where other factors may limit mosquito life cycle Lega et al. (2017).

On the one hand, we did not consider in the model biotic interactions such as prey-predator or food and space competition with other mosquito taxa during the larval stages, despite this is another factor that influences the trajectory of introduced populations (Armistead et al., 2008; Tripet et al., 2011; Reiskind and Lounibos, 2013; Montarsi et al., 2013, 2015b; Müller et al., 2018). On the other hand, we considered the effect of intra-specific competition on larval survival (but not development). We generalised the information available for *Ae. aegypti* to the other *Aedes* species due to the lack of species-specific experiments (Hancock et al., 2016). We recognise that this is not optimal under many facets, but intra-specific larval interactions were a key driver of mosquito-population dynamics that could not be excluded by model structure.

Finally, we did not consider evolutionary processes in our model which may affect invasion success over medium-long time spans. Given the reproductive strategy of *Aedes* mosquitoes, rapid evolutionary processes may take place over relatively short temporal periods (e.g. decades), making introduced populations able to extend their original niche (McBride et al., 2014b).

### 4.2 Proposed research directions

`dynamAedes` is an open-source tool for testing ecological hypothesis and/or to support management plans concerning AIMs. Selecting areas at risk of AIM establishment or periods when abundances are more likely to peak should be considered as facets of AIM surveillance. The importance of such early information becomes fundamental for protecting human health when treating AIM involved in pathogen transmission, as early information on new trajectories of AIM populations becomes critical in the current climate change era. Mosquitoes are affected by temperature changes in, often, predictable ways, though changes in population dynamics can be extremely rapid. Modelling population dynamics under climate change scenarios may thus provide information for anticipating both AIM population changes in space and time and human health risks.

The conceptualisation and design of `dynamAedes` required the review of up-to-date ecological and physiological literature available on four *Aedes* species, which was integrated with feedback from expert ecologists and medical entomologists. It emerged that knowledge on some ecological aspects of these species is highly fragmented or poor (e.g., Cebrián-Camisón et al. 2020), and often dependant on experimental settings and lab strain (e.g., Kramer et al. 2020). Thus, the exploitation of such sets of information for process-based models and, hence, for AIM management would greatly benefit from a standardised review effort and possibly centralised repositories, as already done in other scientific fields such as plant functional traits (Kattge et al., 2020) or systems biology (Tsigkinopoulou et al., 2018). Moreover, experiments on *Ae. japonicus* and *Ae. koreicus* life cycles are just starting to unravel these species eco-physiological rates (*Ae. koreicus*: (Ciocchetta et al., 2017; Marini et al., 2019b); *Ae. japonicus*: (Scott, 2003; Reuss et al., 2018; Wieser et al., 2019) and much work remain to be done on this species. On the contrary, there is large information concerning *Ae. aegypti* and *Ae. albopictus* physiological rates which anyhow have been shown to be highly heterogeneous (Eisen et al., 2014) likely due to non-standardised experimental designs and ecological plasticity of *Aedes* populations (*sensu* Kramer et al. 2021). Information regarding mosquito dispersal is even scarcer and available only for *Ae. aegypti* and *Ae. albopictus*, while passive dispersal through auto-vehicles has been estimated only for *Ae. albopictus* (Eritja et al., 2017) despite the worldwide spread of *Aedes* species was most likely caused by means of passive dispersal. From a biological perspective, future developments of `dynamAedes` may consider also the addition of a `strain` argument, where the physiological temperature-dependent function can be fitted on geographically different mosquito strains, such as tropical, mediter-ranean or temperate (Marini et al., 2020; Kramer et al., 2021). Moreover, if observational data are available, calibration of some parameters values, such as the juvenile density-dependent mortality rate, might be implemented following for instance a Bayesian approach (Marini et al., 2019b).

We believe that a closer interaction between modelers and experimenters will motivate the collection of standardised data on unknown eco-physiological AIM rates that would lead to more accurate model predictions. This project was inspired by such interactions and, in this spirit, `dynamAedes` was meant to be modified or extended to relax its assumptions and limitations with new available information by anyone having basic R programming skills.

## 5 Conclusion

In this study, we presented `dynamAedes`, a mechanistic process-based model to infer invasive *Aedes* mosquito spatio-temporal population dynamics. This first version of the model showed to be often reliable in terms of both biological realism and statistical accuracy. The open-source nature and programming language accessibility and flexibility of the project offers great potential to further develop the model, allowing to better tune the temperature-dependent functions when new physiological observations and findings become available. Abundance estimations derived from `dynamAedes` could be used to inform epidemiological models (e.g., SIR or SEIR) and thus obtain estimations on the risk of pathogens transmission. Finally, it does not seem unrealistic to extend the model application to other species of the genus *Aedes* such as *Ae. notoscriptus* or to species belonging to other genus of medical interest belonging to the *Culicidae* family, such as *Anopheles*, or even to other blood-sucking insects belonging to different taxa such as *Culicoides*.

## 6 Acknowledgements

No conflict of interest has been declared by the author(s).

We thank the French Ministry of Health for providing the confirmed *Ae. albopictus* established population data (SI-LAV), and the EID Mediterranée for providing us the ovitraps data from Nice. Computational resources have been provided by the DART research group at the University of California, Davis as well as supercomputing facilities of the Université catholique de Louvain (CISM/UCL) and the Consortium des Équipements de Calcul Intensif en Fédération Wallonie Bruxelles (CÉCI) funded by the Fond de la Recherche Scientifique de Belgique (F.R.S.-FNRS) under convention 2.5020.11 and by the Walloon Region.

DDR and MM are supported by the FRS-FNRS. MM is also supported by the “Action de Recherche concertée” grant number 17/22-086. FR and RM were supported by the Hessian Centre on Climate Change (FZK) of the Hessian Agency for Nature Conservation, Environment and Geology (HLNUG). Research of RM was funded through the 2018-2019 BiodivERsA joint call for research proposals, under the BiodivERsA3 ERA-Net COFUND programme, and with the funding organisation FWO (BiodivERsA2018-A-323). The Outbreak Research Team of the Institute of Tropical Medicine is financially supported by the Department of Economy, Science and Innovation of the Flemish government.

This project was done within the framework of AIM-COST Action CA17108 (www.aedescost.eu).

## 7 Authors contribution

MM and DDR conceived the ideas, designed the methodology and analysed the data; WVB, FR, RM, SB, FM, SC, DA, GM, AP, GLA, GL and CJMK provided expert opinion on mosquito biology as well as observational datasets to validate the model; DDR, MM and SOV led the writing of the manuscript. All authors contributed critically to the drafts and gave final approval for publication.

## A Review of mechanistic model for invasive *Aedes*

**Table 2:** Description of mechanistic codes for invasive Aedes made available as software or scripts.

| **Name** | **Year** | **Species** | **Biological resolution** | **Time** | **Space** | **Programming language** | **Reference** |
| --- | --- | --- | --- | --- | --- | --- | --- |
| SkeeterBooster | 2009 | *Ae. aegypti* | Agent-based |  |  | C++ | Magori et al. (2009) |
| stagePop | 2009 |  | Process-based |  |  | R | Kettle and Nutter (2015) |
| sPop | 2018 |  |  |  |  | C++, R, Python | Erguler (2018) |
| albopictus | 2019 | *Ae. albopictus* | Process-based |  |  | C++, R, Python 4 | Erguler et al. (2016, 2017) |
|  | 2019 | *Ae. albopictus* | Process-based |  |  | Octave v4.2.1, Runge-Kutta 4 | Metelmann et al. (2019) |
|  | 2019 | *Ae. albopictus* | Process-based |  |  | R, Ocelet | Tran et al. (2013, 2020) |
|  | 2020 | *Ae. aegypti* | Process-based |  |  | R | Iwamura et al. (2020) |
|  | 2020 | *Ae. koreicus* | Process-based |  |  | R | Kurucz et al. (2020) |

## B Parameters used in dynamAedes

**Table 3:**
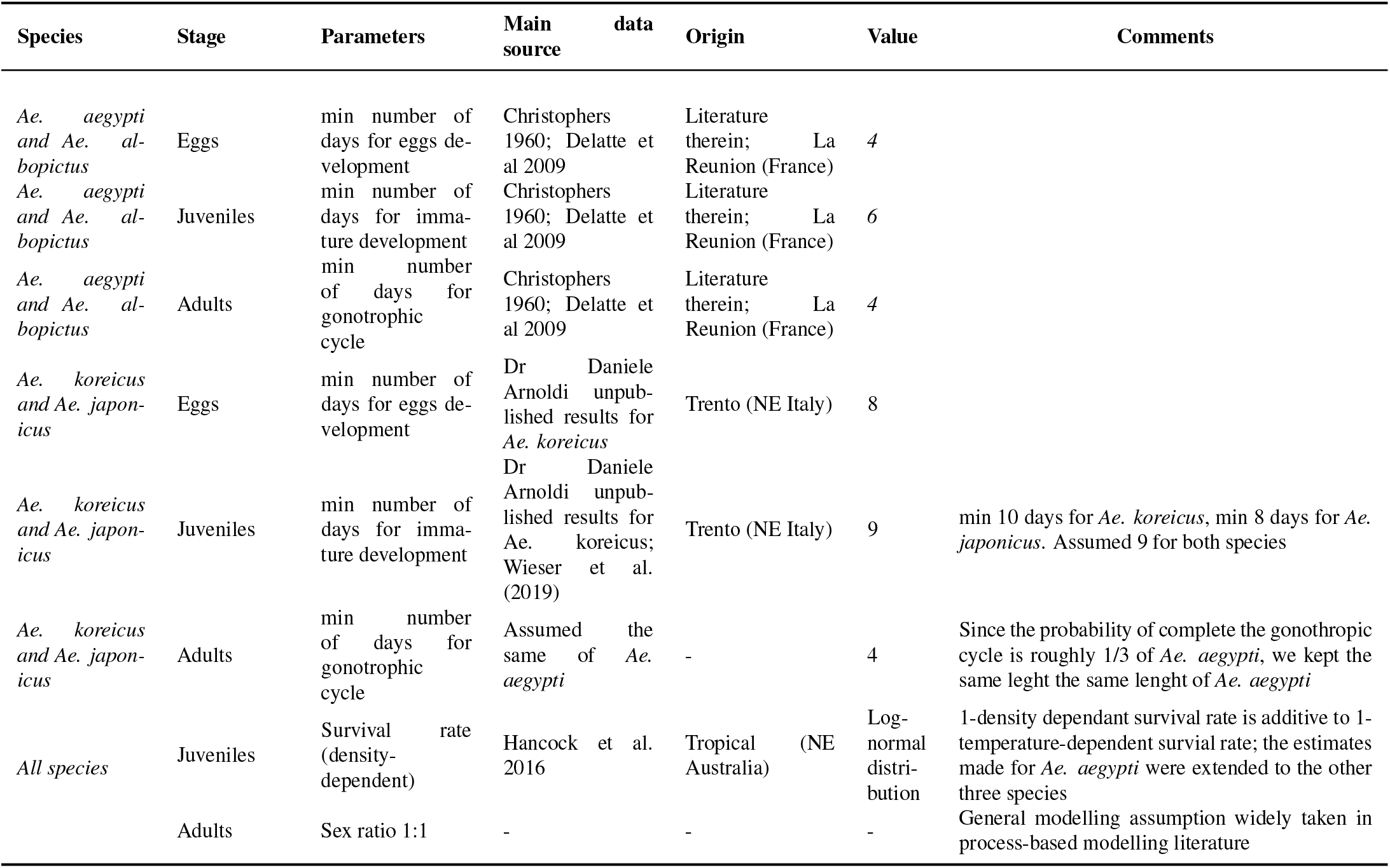
Other model features

**Table 4:**
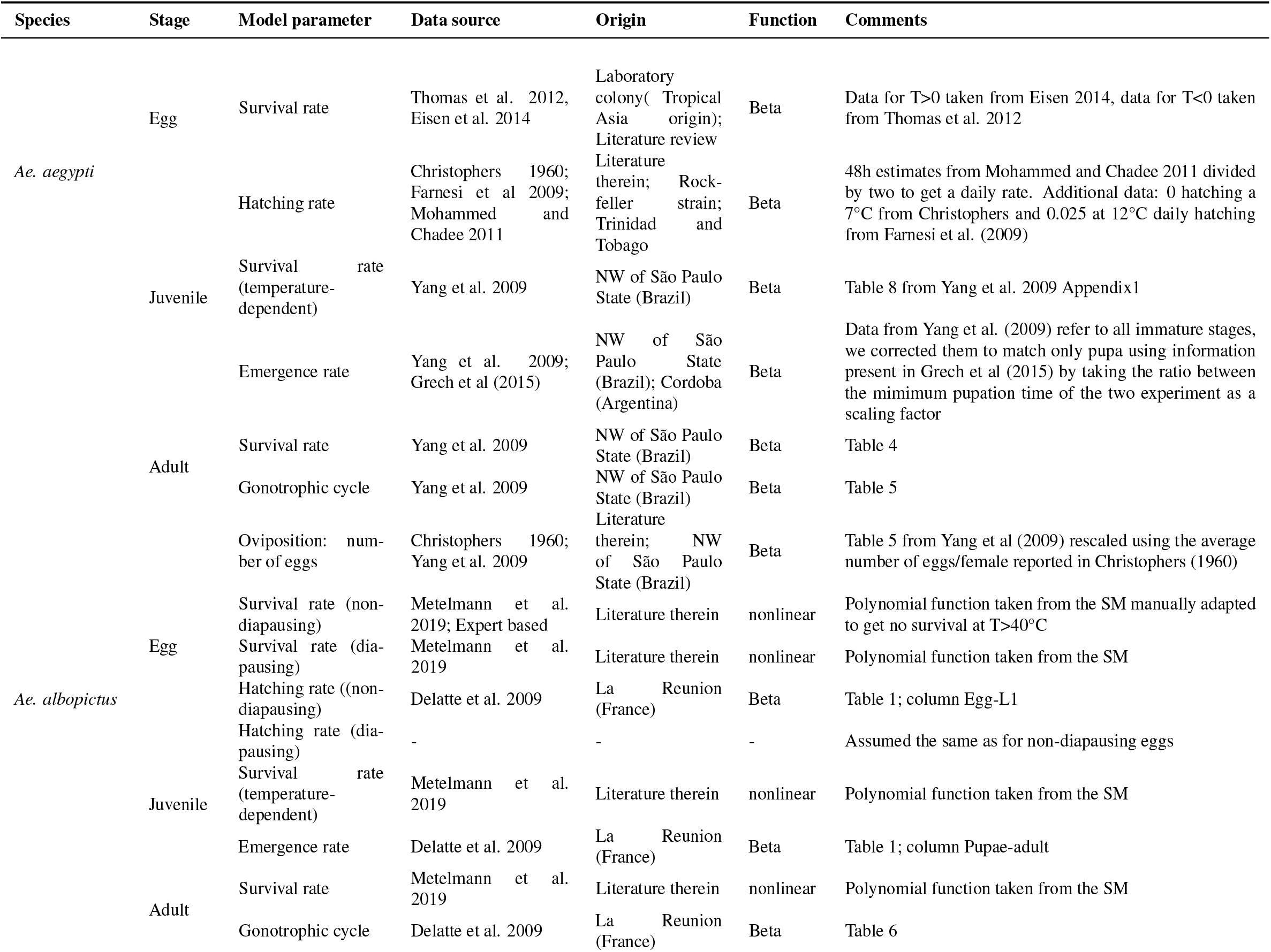
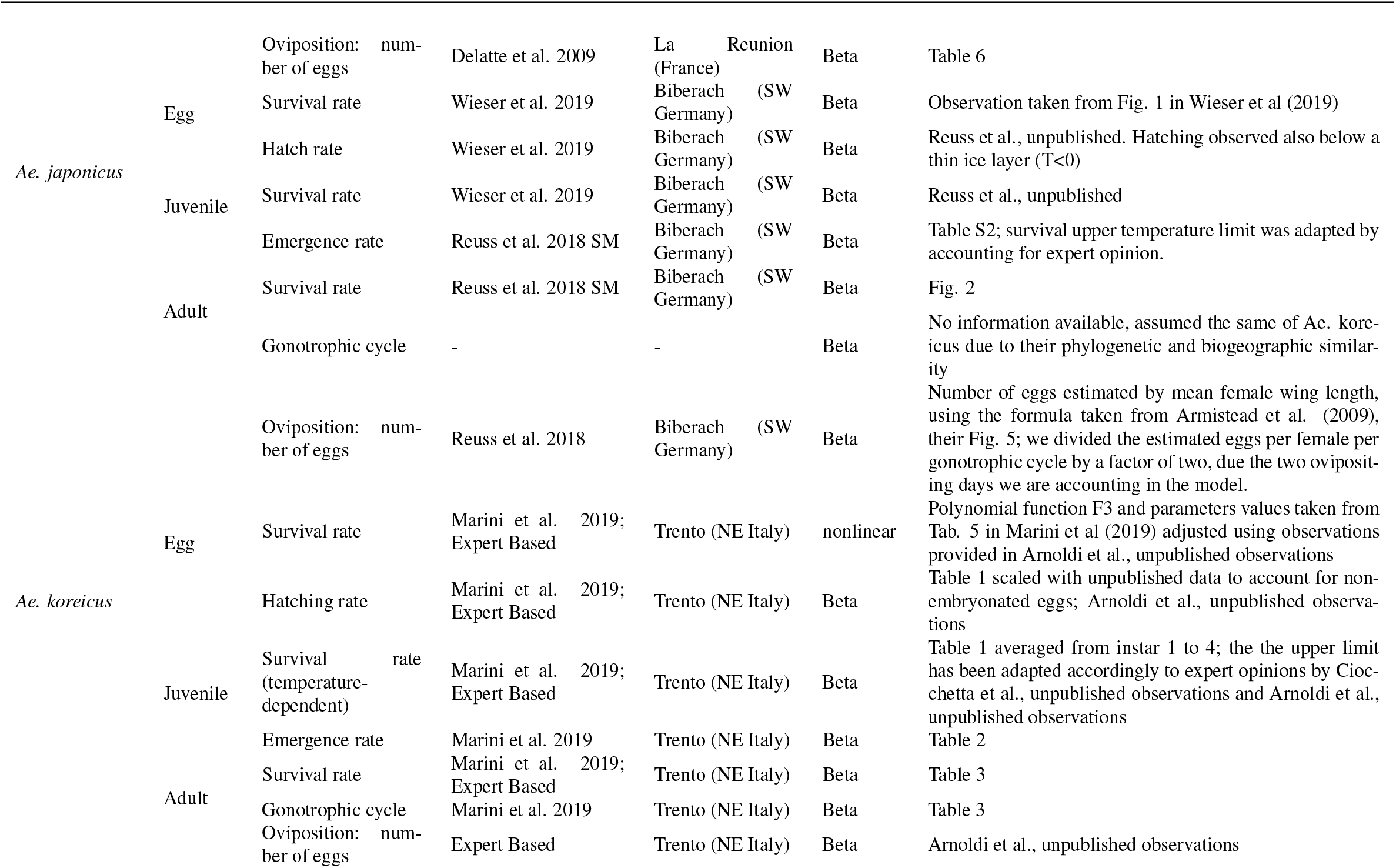
Species-specific temperature-dependent physiological parameters

**Table 5:** Species specific dispersal parameters

| Species | Stage | Parameters | Main source | Origin | Distribution | Comments |
| --- | --- | --- | --- | --- | --- | --- |
| *Ae. aegypti* | Adults | Active dispersal | Marcantonio et al. 2019 | California (USA) | Log-normal | Table 3 |
| *Ae. albopictus* | Adults | Active dispersal | Marini et al. 2019B | Rome (Italy) | Log-normal | Figure 2a; Data from the 3 MRR were fit with several distributions, best distribution chosen using AIC |
| *Ae. japonicus* | Adults | Active dispersal | - | - | Log-normal | No information available, assumed the same of *Ae. albopictus* |
| *Ae. koreicus* | Adults | Active dispersal | - | - | Log-normal | No information available, assumed the same of *Ae. albopictus* |
| All species | All species | Passive dispersal (average trip distance) | Pasaoglu et al. 2012 | - | - | Trip distance weighted average for ITA, DEU, FRA, ESP, POL, UK. Data taken from Figure 4.4 Average trip distance (km) by trip purpose using WebPlotDigitalizer |
| All species | Adults | Passive dispersal (Hitchhiking probability) | Ertijia et al. 2017 | Mediterranean (Catalonia, Spain) | gamma distribution | 0.0051 probability of a female mosquito to enter a car; Assumed to be the same for all the species as estimated by Ertijia et al. 2017 for *Ae. albopictus* |

**Table 6:**
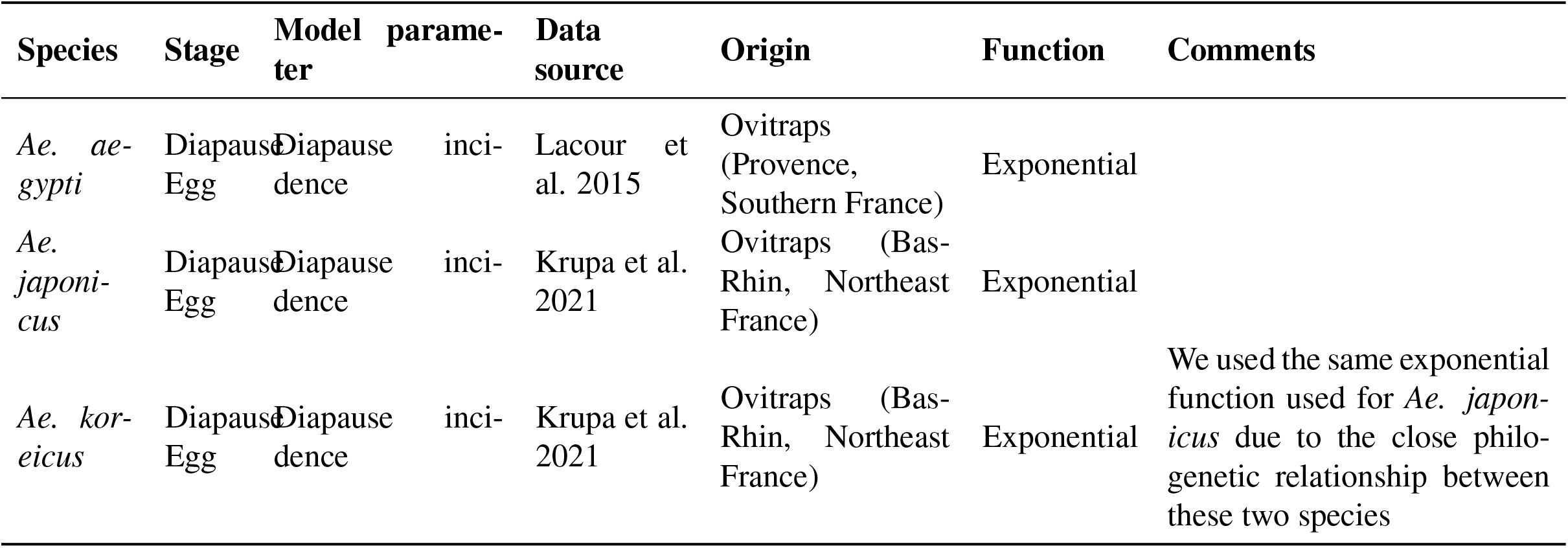
Species specific photoperiod parameters

| Species | Stage | Model parameter | Data source | Origin | Function | Comments |
| --- | --- | --- | --- | --- | --- | --- |
| *Ae. aegypti* | Diapause Egg | Diapause incidence | Lacour et al. 2015 | Ovitrap (Provence, Southern France) | Exponential |  |
| *Ae. japonicus* | Diapause Egg | Diapause incidence | Krupa et al. 2021 | Ovitrap (Bas-Rhin, Northeast France) | Exponential |  |
| *Ae. kor-eicus* | Diapause Egg | Diapause incidence | Krupa et al. 2021 | Ovitrap (Bas-Rhin, Northeast France) | Exponential | We used the same exponential function used for *Ae. japonicus* due to the close phylogenetic relationship between these two species |

## A *Aedes sp.* response curve

**Figure 7:**
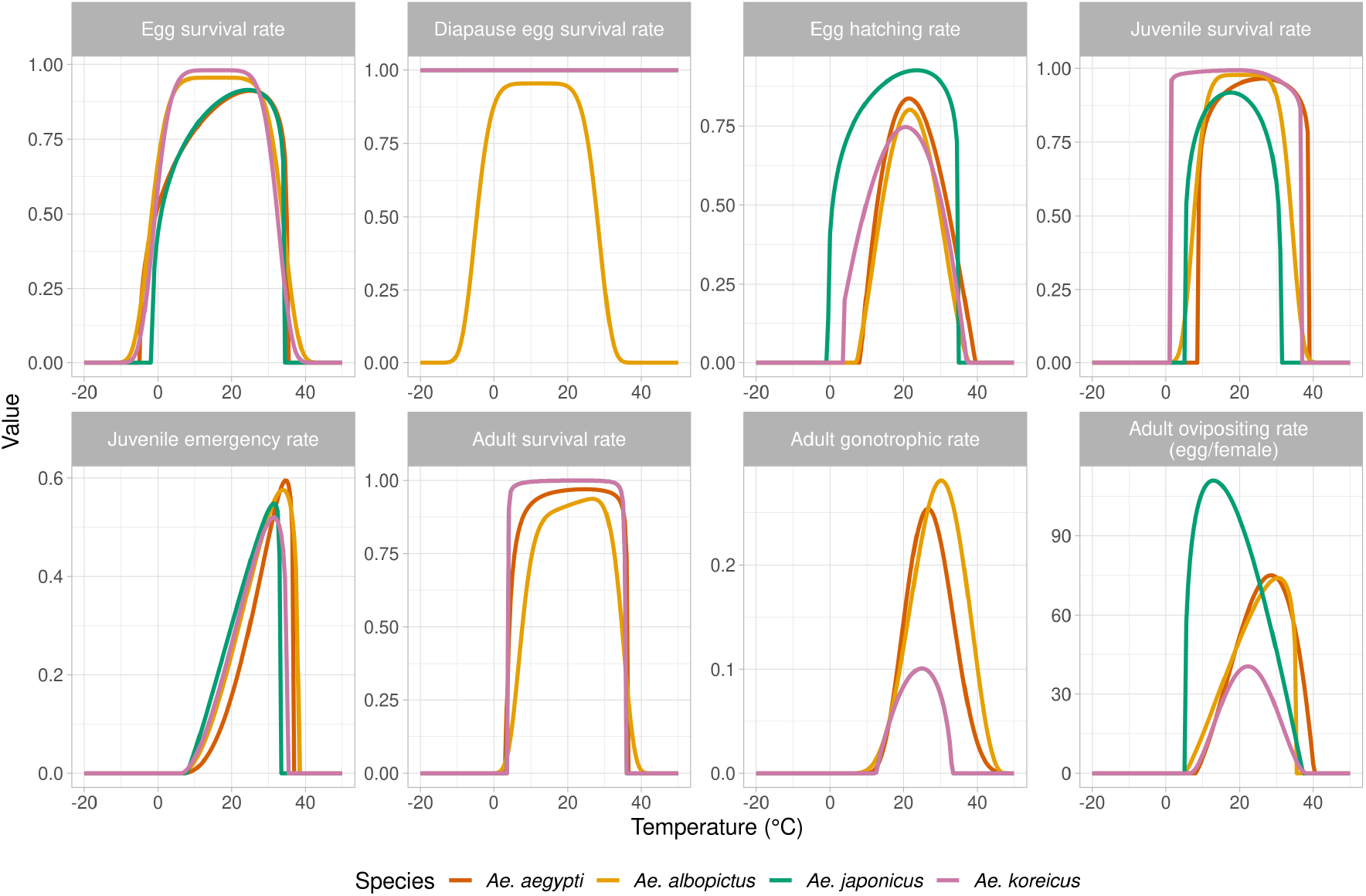
Overview of the temperature-dependent functions used in the model for the four *Aedes species*

**Figure 8:**
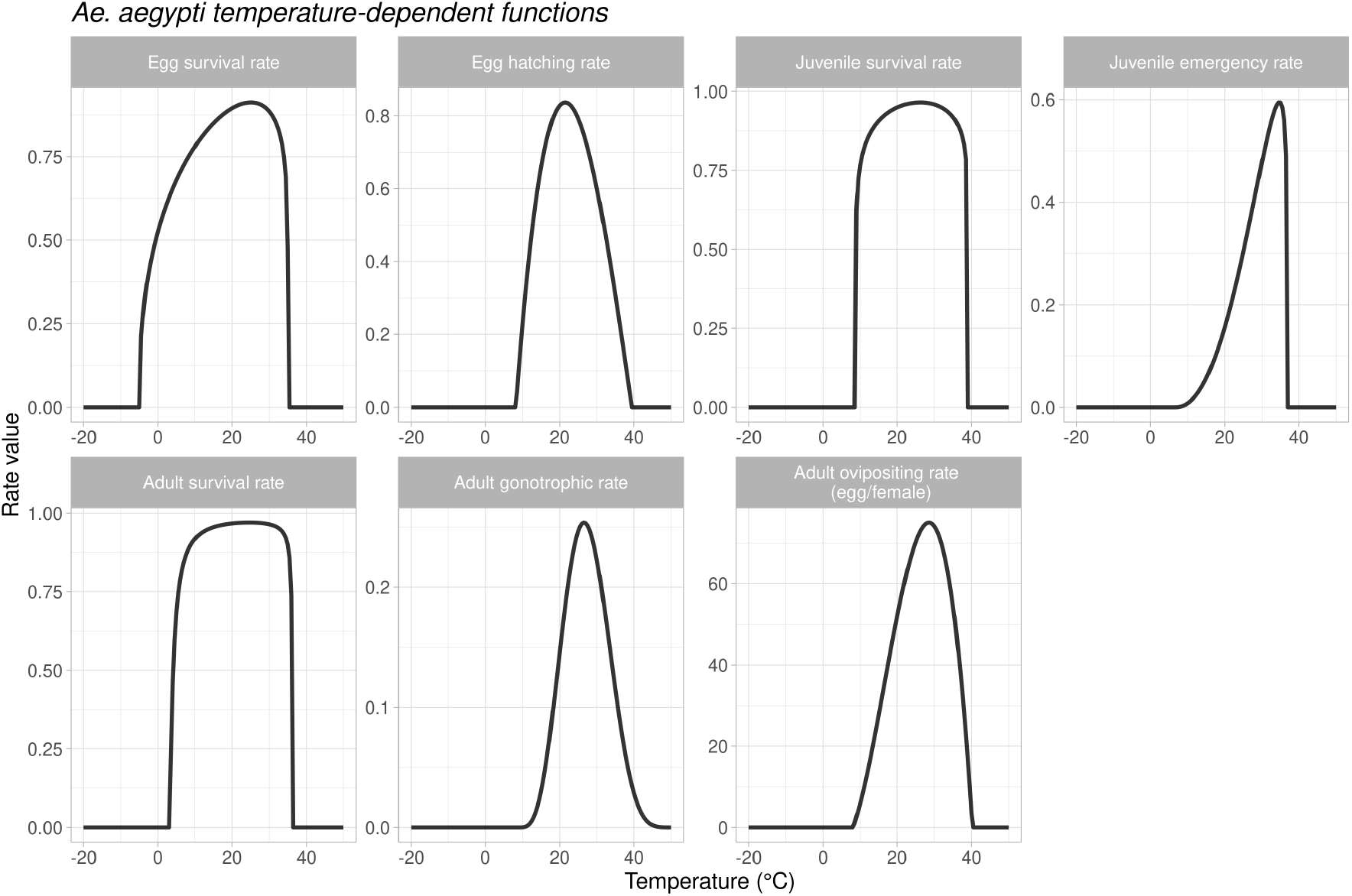
Overview of the temperature-dependent functions used in the model for *Ae. aegypti*

**Figure 9:**
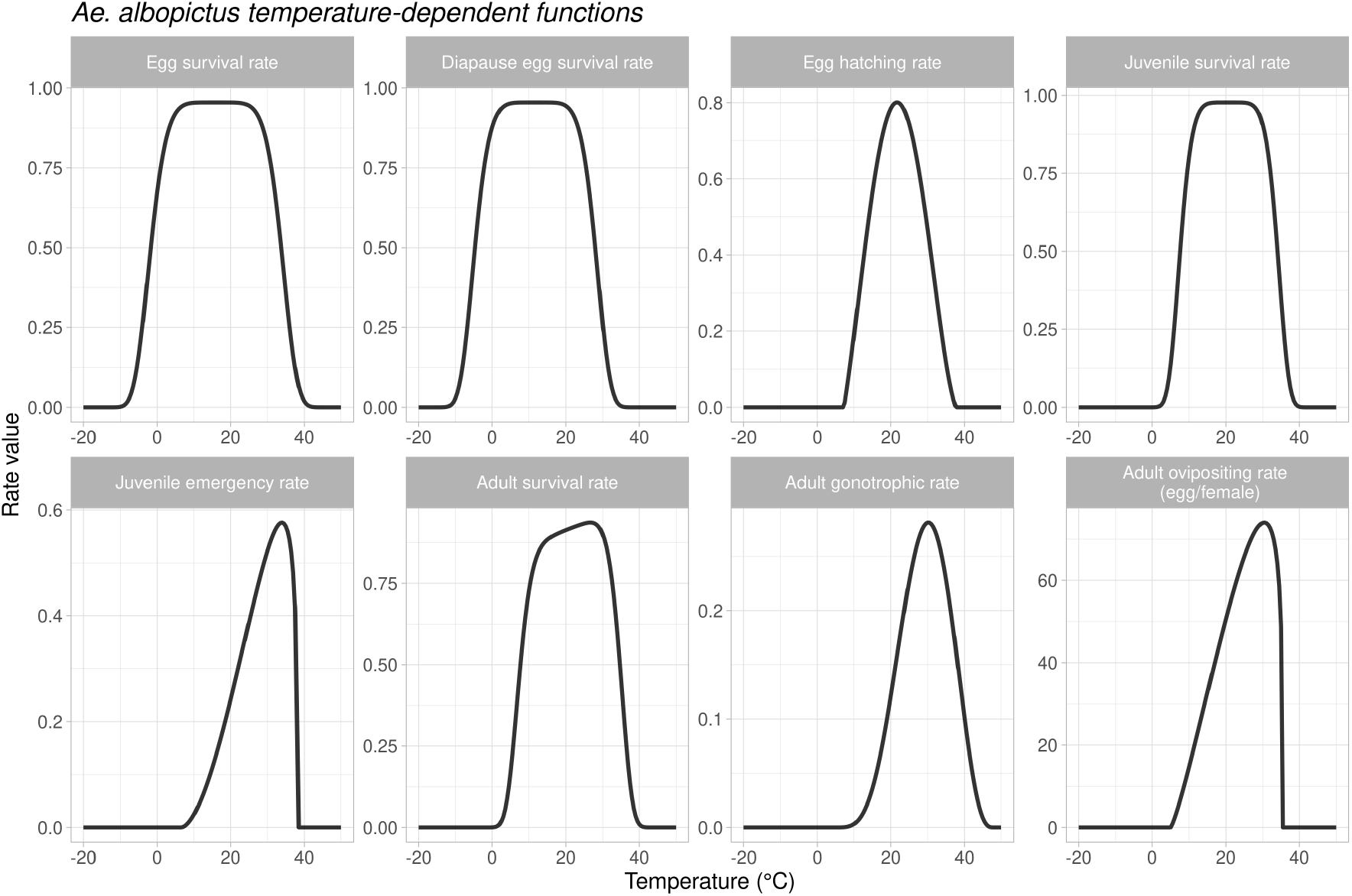
Overview of the temperature-dependent functions used in the model for *Ae. albopictus*

**Figure 10:**
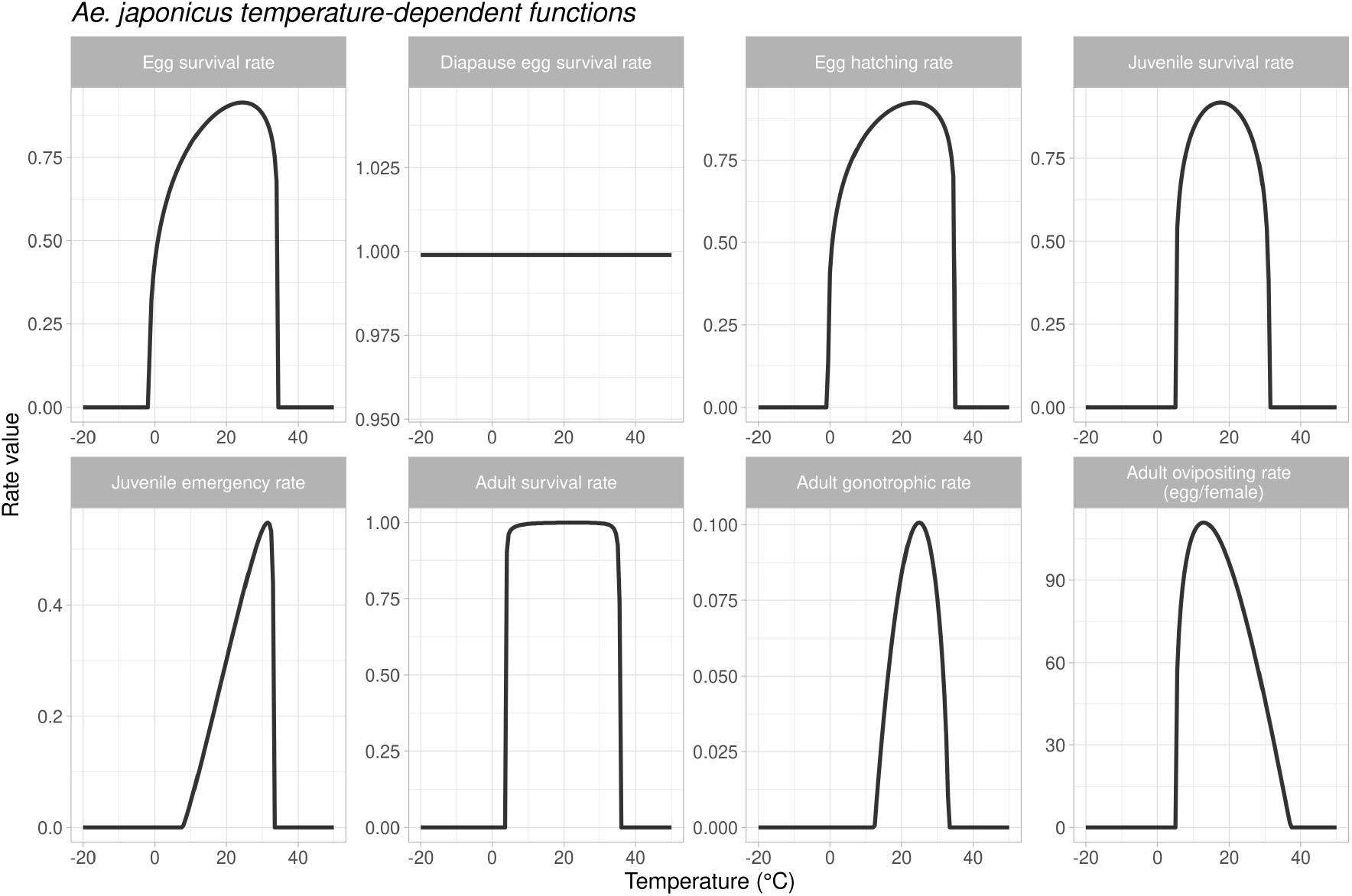
Overview of the temperature-dependent functions used in the model for *Ae. japonicus*

**Figure 11:**
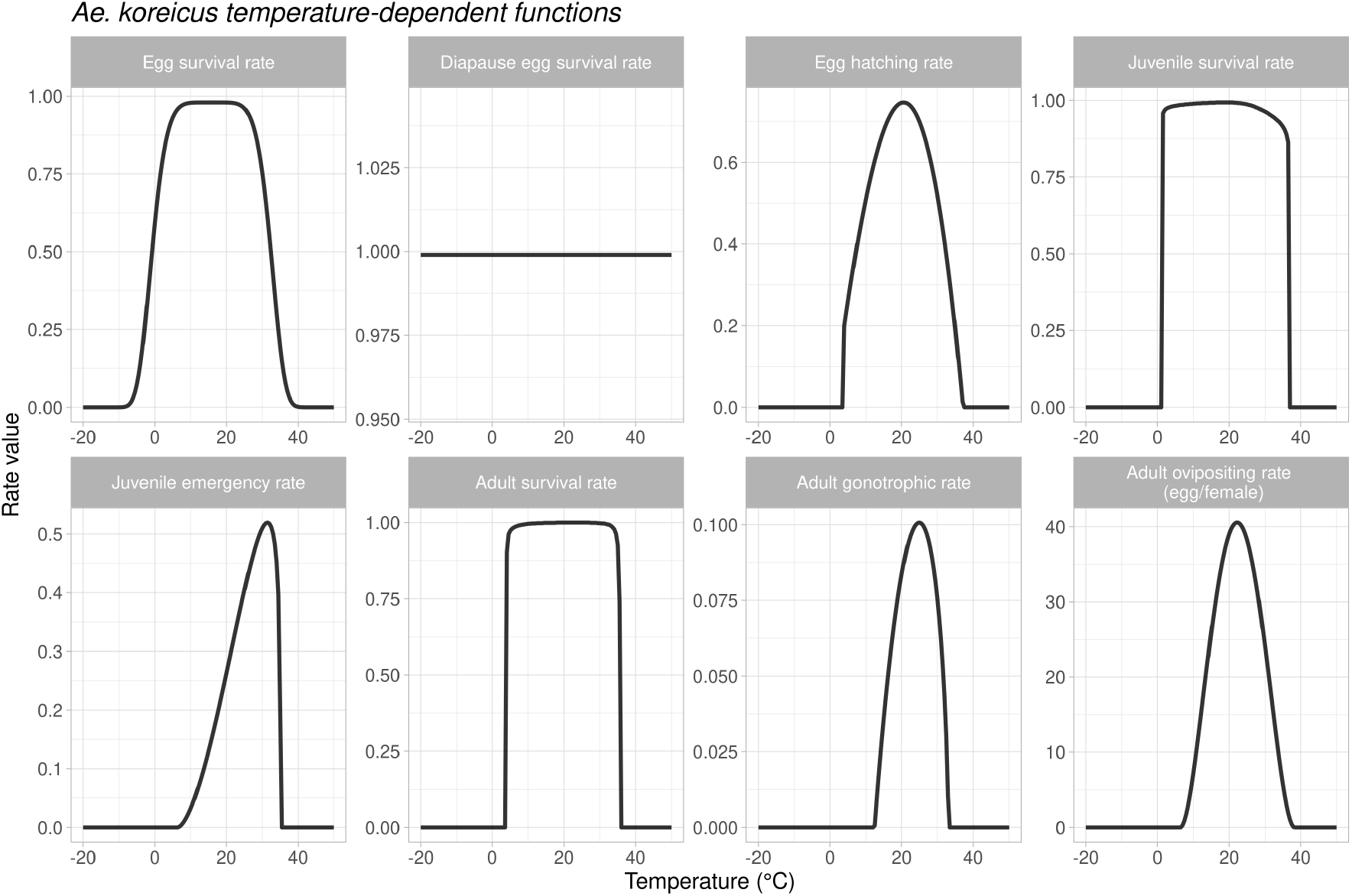
Overview of the temperature-dependent functions used in the model for *Ae. koreicus*

**Figure 12:**
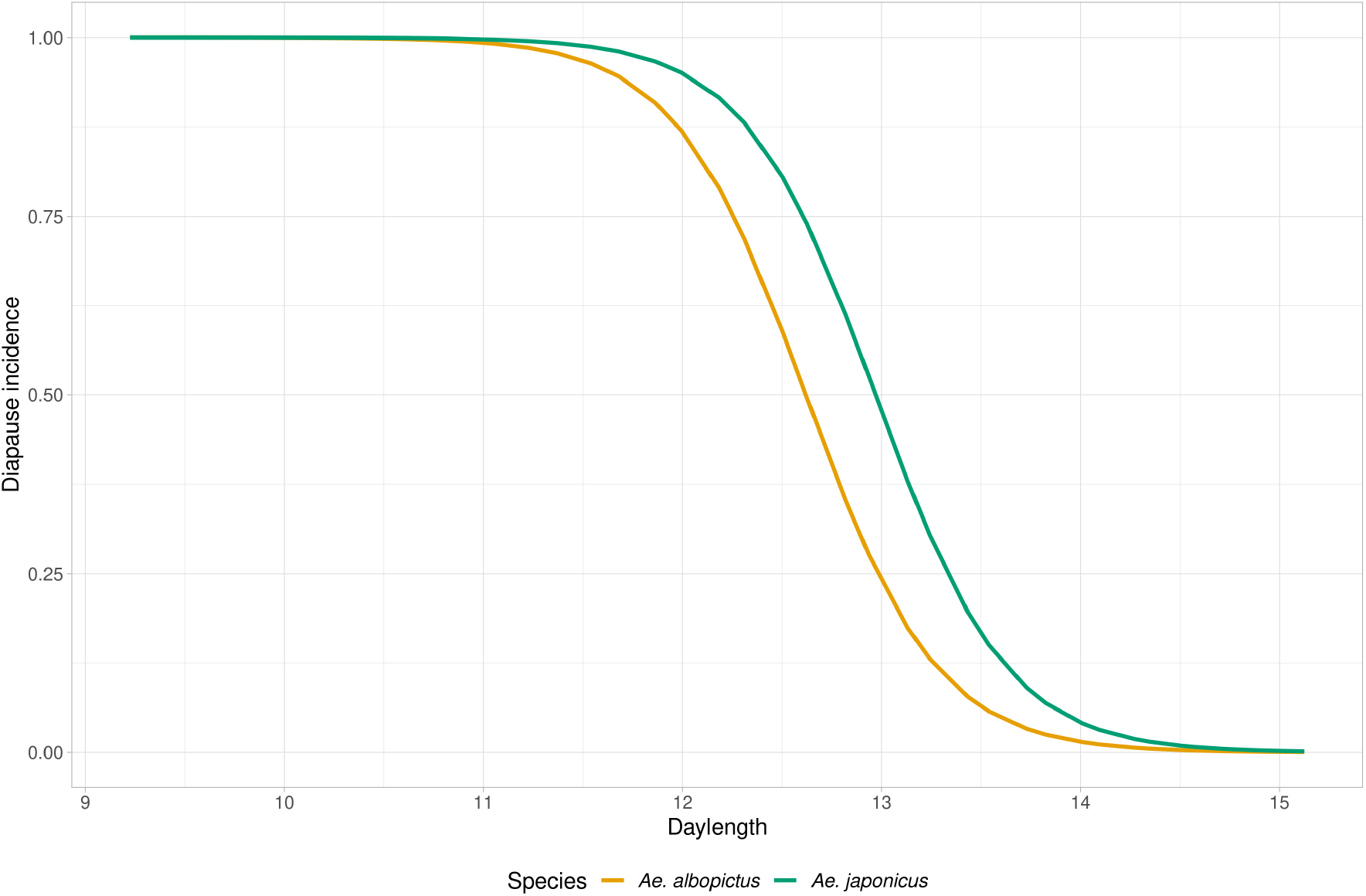
Overview of the photoperiod-dependent diapause incidence function used to in the model for *Ae. albopictus* and *Ae. japonicus*. The *Ae. japonicus* function was used for *Ae. koreicus* as well.

## B Larval habitat water volume parameter sensitivity

**Figure 13:**
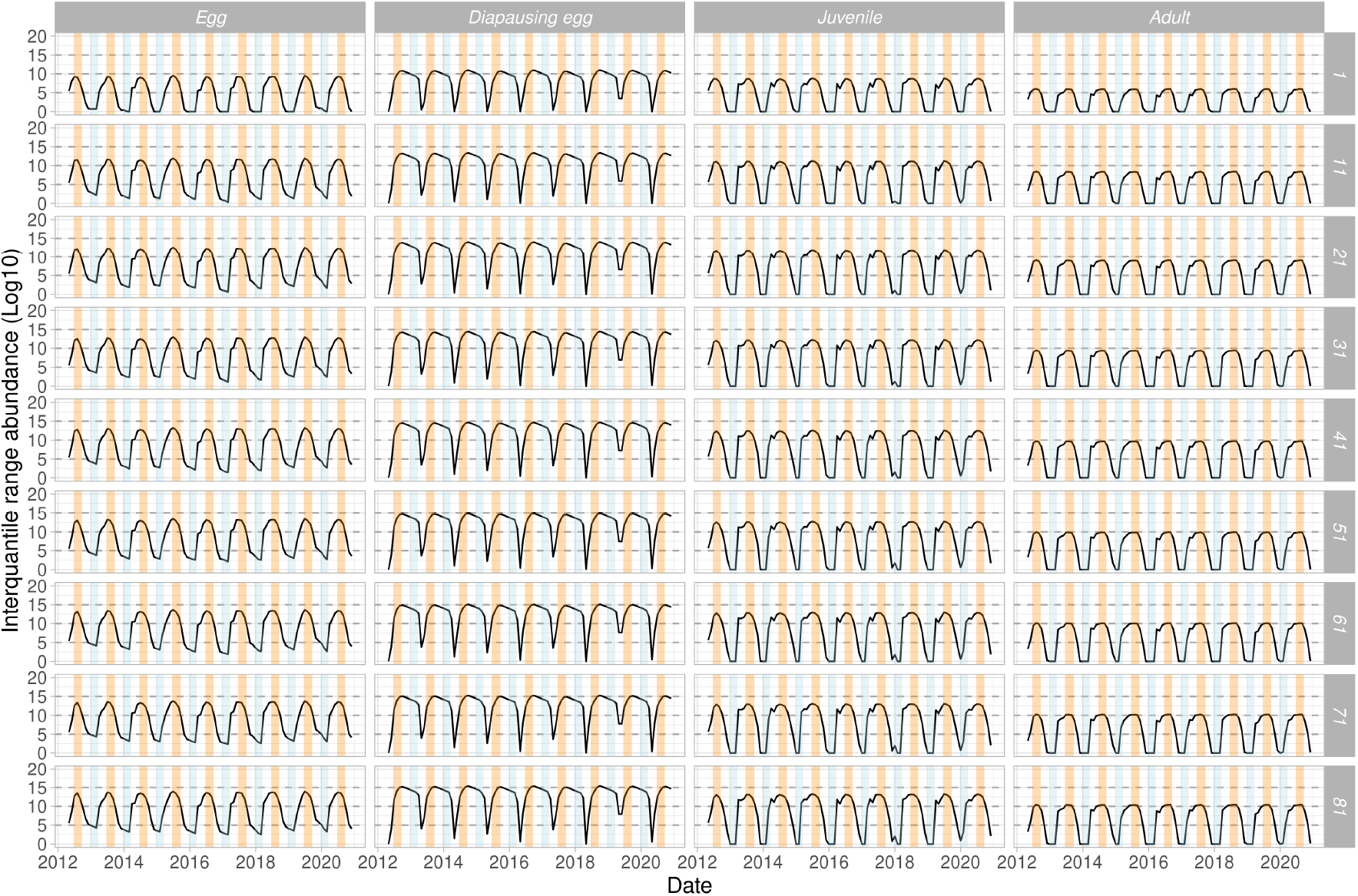
Sensitivity analysis on the effect of the `lhwv` parameter on the estimated number of individuals. In this example, we used the temperature observations of the Nice weather station used in the case study and varied the amount of water volume. The seasonal trend remained the same but, as expected, the simulated number of individuals increase as the `lhwv` parameter increase.

## C *Aedes aegypti* and *Ae. albopictus* regional scale case study

**Figure 14:**
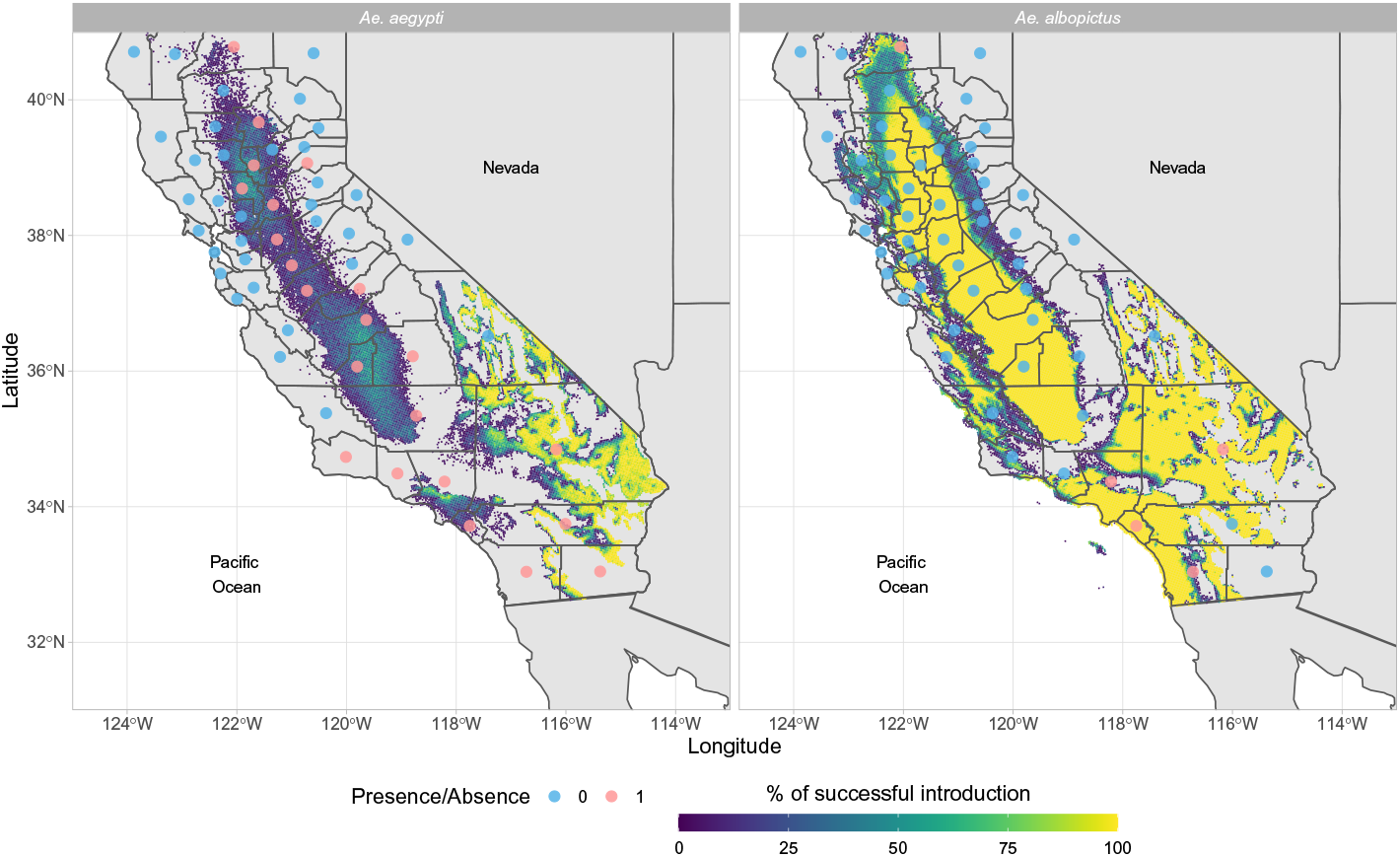
Predicted percentage of establishment of *Ae. aegypti Ae. albopictus* in California (USA) for the years 2011-2016 and 2013-2018, respectively. Only pixels having a probability of successful introduction >0 are shown. The red dots represent the counties where the species have been found.

## D *Aedes koreicus* population dynamics punctual scale case study

**Table 7:** Model validation for *Aedes koreicus* model in Trento (NE Italy)

| Year | Month | CI 2.5% | CI 25% | CI 50% | CI 75% | CI 97.5% | Observed *Ae. Koreicus* |
| --- | --- | --- | --- | --- | --- | --- | --- |
| 2016 | May | 0.00 | 1.45 | 5.26 | 13.74 | 31.22 | 2 |
| 2016 | June | 0.00 | 1.37 | 6.36 | 14.40 | 40.64 | 5 |
| 2016 | July | 0.00 | 2.32 | 11.85 | 32.62 | 101.76 | 3 |
| 2016 | August | 0.70 | 24.57 | 66.65 | 128.03 | 266.00 | 9 |
| 2016 | September | 0.18 | 14.52 | 37.37 | 88.27 | 175.66 | 37 |
| 2016 | October | 0.00 | 0.24 | 1.18 | 3.14 | 10.17 | 0 |
| 2017 | May | 0.02 | 0.51 | 2.67 | 7.85 | 18.44 | 9 |
| 2017 | June | 0.29 | 2.75 | 10.60 | 28.57 | 55.28 | 8 |
| 2017 | July | 3.67 | 10.91 | 30.62 | 85.53 | 186.93 | 10 |
| 2017 | August | 7.24 | 32.50 | 74.18 | 159.67 | 283.74 | 24 |
| 2017 | September | 0.61 | 8.79 | 31.48 | 88.08 | 156.47 | 2 |
| 2017 | October | 0.08 | 0.82 | 2.20 | 4.98 | 11.06 | 2 |
| 2017 | November | 0.00 | 0.00 | 0.00 | 0.00 | 0.14 | 0 |
| 2018 | April | 1.66 | 5.26 | 11.54 | 22.25 | 34.30 | 0 |
| 2018 | May | 1.73 | 3.22 | 5.73 | 9.18 | 13.21 | 1 |
| 2018 | June | 1.57 | 5.81 | 10.52 | 22.29 | 32.89 | 6 |
| 2018 | July | 13.74 | 29.99 | 48.04 | 74.34 | 98.01 | 7 |
| 2018 | August | 37.83 | 71.67 | 109.27 | 157.55 | 201.00 | 29 |
| 2018 | September | 54.55 | 82.03 | 112.57 | 147.58 | 179.09 | 24 |
| 2018 | October | 2.13 | 2.90 | 3.77 | 5.89 | 7.80 | 6 |
| 2018 | November | 0.00 | 0.00 | 0.00 | 0.08 | 0.15 | 0 |

## Footnotes

1 AedesRisk v1.0: https://shinyapps.ecdc.europa.eu/shiny/AedesRisk/

2 See https://maps.vectorsurv.org/

3 California Department of Public Health, “Map and City List of *Aedes aegypti* and *Aedes albopictus* Mosquitoes in CA, 2011-2021” (accessed on 28th October 2021): https://www.cdph.ca.gov/Programs/CID/DCDC/Pages/VBDS.aspx

4 CDS Toolbox: https://cds.climate.copernicus.eu/toolbox/doc/index.html

5 https://signalement-moustique.anses.fr/signalement_albopictus/

6 https://www.data.gouv.fr/fr/datasets/trafic-moyen-journalier-annuel-sur-le-reseau-routier-national/

7 https://www.leparisien.fr/economie/l-europe-fait-passer-la-lgv-paris-lyon-a-l-heure-php

8 https://solidarites-sante.gouv.fr/sante-et-environnement/risques-microbiologiques-physiques-et-chimiques/especes-nuisibles-et-parasites/article/cartes-de-presence-du-moustique-tigre-aedes-albopictus-en-france-metropolitain

9 www.meteotrentino.it

## Notes

### Competing Interest Statement

The authors have declared no competing interest.

